# The ESCRT protein CHMP5 promotes T cell leukemia by controlling BRD4-p300-dependent transcription

**DOI:** 10.1101/2024.01.29.577409

**Authors:** Katharine Umphred-Wilson, Shashikala Ratnayake, Qianzi Tang, Rui Wang, Ballachanda N. Devaiah, Lan Zhou, Qingrong Chen, Daoud Meerzaman, Dinah S Singer, Stanley Adoro

**Affiliations:** Experimental Immunology Branch, National Cancer Institute, National Institutes of Health, Bethesda, MD 20892; Immunology Training Program, Department of Pathology, Case Western Reserve University School of Medicine, Cleveland, OH 44106, USA; Computational Genomics and Bioinformatics Branch, Center for Biomedical Informatics & Information Technology, National Cancer Institute, National Institutes of Health, Bethesda, MD 20850; College of Animal Science and Technology, Sichuan Agricultural University; Chengdu 611130, China; Department of Pathology and Genomic Medicine, Houston Methodist Hospital, Houston, TX 77030

## Abstract

Oncogene activity rewires cellular transcription, creating new transcription networks to which cancer cells become addicted, by mechanisms that are still poorly understood. Using human and mouse models of T cell acute lymphoblastic leukemia (T-ALL), we identify an essential nuclear role for CHMP5, a cytoplasmic endosomal sorting complex required for transport (ESCRT) protein, in establishing and maintaining the T-ALL transcriptional program. Nuclear CHMP5 promoted the T-ALL gene program by augmenting recruitment of the co-activator BRD4 by the histone acetyl transferase p300 selectively at enhancers and super-enhancers, an interaction that potentiated H3K27 acetylation at these regulatory enhancers. Consequently, loss of CHMP5 diminished BRD4 occupancy at enhancers and super-enhancers and impaired RNA polymerase II pause release, which resulted in downregulation of key T-ALL genes, notably *MYC*. Reinforcing its importance in T-ALL pathogenesis, CHMP5 deficiency mitigated chemoresistance in human T-ALL cells and abrogated T-ALL induction by oncogenic NOTCH1 *in vivo*. Thus, the ESCRT protein CHMP5 is an essential positive regulator of the transcriptional machinery promoting T-ALL disease.

**Graphical abstract:** 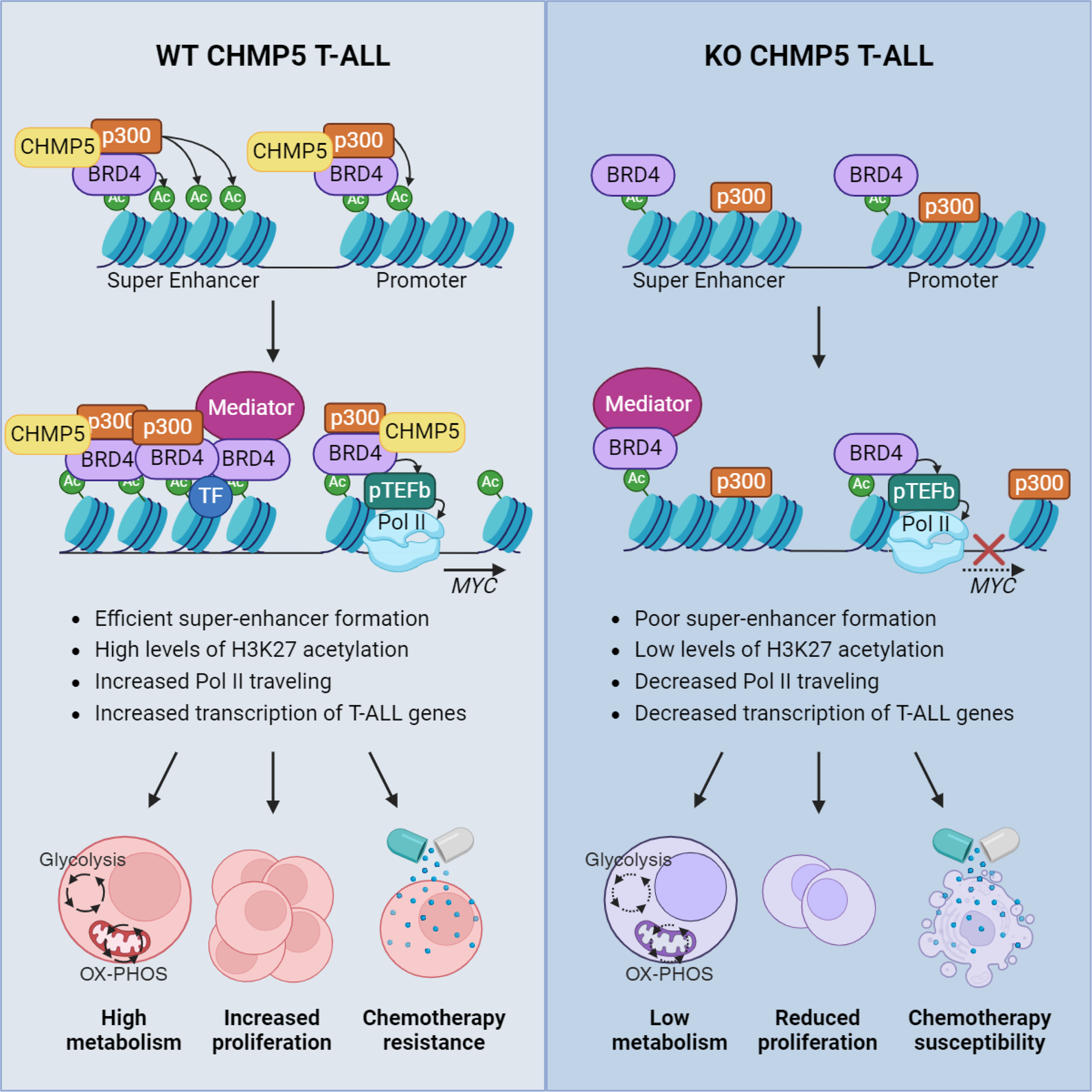

**Highlights:**

- Identification of a nuclear role for the cytosolic ESCRT protein CHMP5 in transcription
- CHMP5 mediates BRD4-dependent Pol II pause release and transcription of T-ALL genes
- P300-BRD4 induced enhancer and super-enhancer H3K27 acetylation requires CHMP5
- CHMP5 depletion mitigates chemoresistance and abrogates T-ALL initiation *in vivo*

## Introduction

Transcriptional dysregulation is a hallmark of cancer cells as oncogenes hijack the transcriptional machinery to promote expression of genes that support their survival and proliferative needs. As “addiction” to the rewired transcriptional machinery is necessary for tumor progression, mediators of transcriptional addiction have emerged as attractive targets against cancers^1,2^. Thus, discovering critical components of the dysregulated transcriptional machinery is essential to understanding cancer biology, identifying therapeutic targets, and improving cancer cell response to drugs. Studies in solid tumors and hematological cancers implicate a crucial role for the bromo and extraterminal (BET) domain protein BRD4 in promoting the cancer gene program especially in MYC-dependent tumors^3–6^. BRD4 is a pleiotropic transcription factor, with a histone acetyltransferase activity via which it acetylates histones to promote nucleosome decompaction, and a kinase activity by which it can phosphorylate and induce RNA polymerase II (Pol II) pause release and transcription^7–11^. In BRD4-dependent cancer cells, key pro-tumorigenic genes display marked enrichment for BRD4 at enhancers and super-enhancers, DNA regions defined by histone hyperacetylation modifications, including histone 3 lysine 27 acetylation (H3K27ac)^12,13^. The presence and dependency of cancer gene expression on these hyperacetylated enhancers distinguishes normal from cancer cells^1,2^, and raises the fundamental question how oncogenes, utilizing the same transcription factors as normal cells, selectively modulate only specific regulatory elements in the cancer cell genome.

T cell acute lymphoblastic leukemia (T-ALL) is an aggressive hematological malignancy characterized by aberrant development and proliferation of immature thymocytes which populate the peripheral circulation and infiltrate vital organs^14^. More than half of all human T-ALL cases are caused by activating NOTCH1 mutations that constitutively generate the intracellular NOTCH1 domain (ICN1)^15^, a transcription factor that induces the proto-oncogene MYC to which many T-ALL cells become dependent^16^. Importantly, ICN1-driven T-ALL exemplifies a cancer with high BRD4 dependency wherein BRD4, cooperatively with ICN1 and MYC, drive expression of key T-ALL genes in a positive feed-forward loop to establish the leukemogenic program^17–19^. Accordingly, BET inhibitors which release BRD4 from chromatin downregulate expression of T-ALL genes like *MYC* and, alone or in combinations with other chemotherapy drugs, suppress T-ALL cell survival *in vitro* and in pre-clinical models^5,17,20^. In addition, BRD4-dependent mechanisms, characterized by enriched BRD4 occupancy at super-enhancers of T-ALL genes, have also emerged as an underlying driver of chemoresistance in T-ALL cells^20,21^. Therefore, elucidating cellular factors that modulate BRD4 recruitment and interaction with chromatin and chromatin modifying factors can uncover novel insights into chemoresistance mechanisms in T-ALL.

Charged multivesicular body protein-5 (CHMP5 or VPS60) is a ∼35 kDa coiled-coil protein first identified in yeast^22,23^ as part of the cytosolic endosomal sorting complex required for transport (ESCRT)-III machinery where it promotes activation of the AAA-ATPase VPS4^24,25^. Because its function in membrane remodeling is required for many cellular processes, the ESCRT machinery has long been linked to cancer, but the precise role of individual ESCRT proteins in tumorigenesis remains unclear and often attributed to their membrane scission activity^26–28^. However, recent studies in osteoclasts and thymocytes, revealed CHMP5 as an adaptor for deubiquitylating enzymes whereby it promoted the stabilization of client proteins required for differentiation and survival^29,30^. Intriguingly, in these hematopoietic cell lineages, ESCRT-mediated processes (including membrane receptor recycling, multivesicular body formation) were not impaired by CHMP5 deficiency^29,30^. This not only suggested a redundant role for CHMP5 in the ESCRT machinery but also indicated evolved CHMP5 functions beyond the ESCRT machinery. Yet, non-membrane remodeling roles of CHMP5 in either normal or cancer cells remain largely unexplored.

T-ALL oncogenes usually co-opt the same cellular factors that promote normal thymocyte development to establish leukemia^31^. Despite its requirement for normal thymocyte development^30^, it is not known whether T-ALL development requires CHMP5. We now report a novel nuclear function for CHMP5 required for T-ALL pathogenesis. Nuclear CHMP5 transcriptionally promoted T-ALL initiation and maintenance by facilitating the recruitment of BRD4 by the histone acetyl transferase p300, an interaction that potentiated H3K27 acetylation selectively at the enhancers and super-enhancers that drive high transcription of key T-ALL genes. Consequently, CHMP5 depletion impaired T-ALL maintenance in vitro and abrogated oncogenic NOTCH1-initiated T-ALL in mice.

## Results

### CHMP5 enables expression of a T-ALL gene program exemplified by MYC

As more than half of human T-ALL cases have underlying NOTCH1 activating mutations that generate ICN1^15^, we set out to define CHMP5 dependent processes and function in T-ALL cells using the CUTLL1 cell line, a CD4^+^CD8^+^ human T-ALL subtype caused by activated NOTCH1^32^. Specifically, CUTLL1 cells harbor a t(7;9) (q34;q34) translocation of *NOTCH1* into the *TCRB* loci that results in constitutive cleavage of NOTCH1 into ICN1 by γ-secretase^32^. We depleted CHMP5 by short hairpins RNA (shRNA) and compared shCHMP5 knockdown (KD) cells to non-targeting control shRNA (CT) cells (**Figure S1A**). CHMP5 depletion had no impact on T-ALL cell viability (**Figure S1B**), but EdU (5-ethynyl-2’-deoxyuridine) incorporation indicated a proliferation defect in KD cells which appeared arrested at S phase with impaired G2/M progression (**Figure S1C**).

To determine the molecular consequences of CHMP5 deficiency in these T-ALL cells, we subjected them to RNA-seq. Comparison of transcriptomes of KD to control T-ALL cells revealed substantial gene expression changes (fold-change ≥ 1.2; adjusted p < 0.05) due to loss of CHMP5; differentially expressed genes (DEG) included 1057 upregulated and 702 downregulated genes. Unbiased pathway analysis of these DEGs revealed significant (p < 0.05; *FDR* < 0.1) upregulation of pathways associated with impaired leukemogenesis including P53, apoptosis, interferon signaling (**Figure 1A**). In parallel, CHMP5-deficient T-ALL cells displayed a striking downregulation of “MYC target” genes that included several MYC-regulated genes involved in glycolysis, oxidative phosphorylation, and cell cycle progression (**Figures 1A, 1B** and **S1D**). Given the essential role of MYC in NOTCH1-driven T-ALL^16–19^, these data suggest a role for CHMP5 in promoting the core transcriptional program induced by oncogenic NOTCH1 in T-ALL.

**Figure 1.**
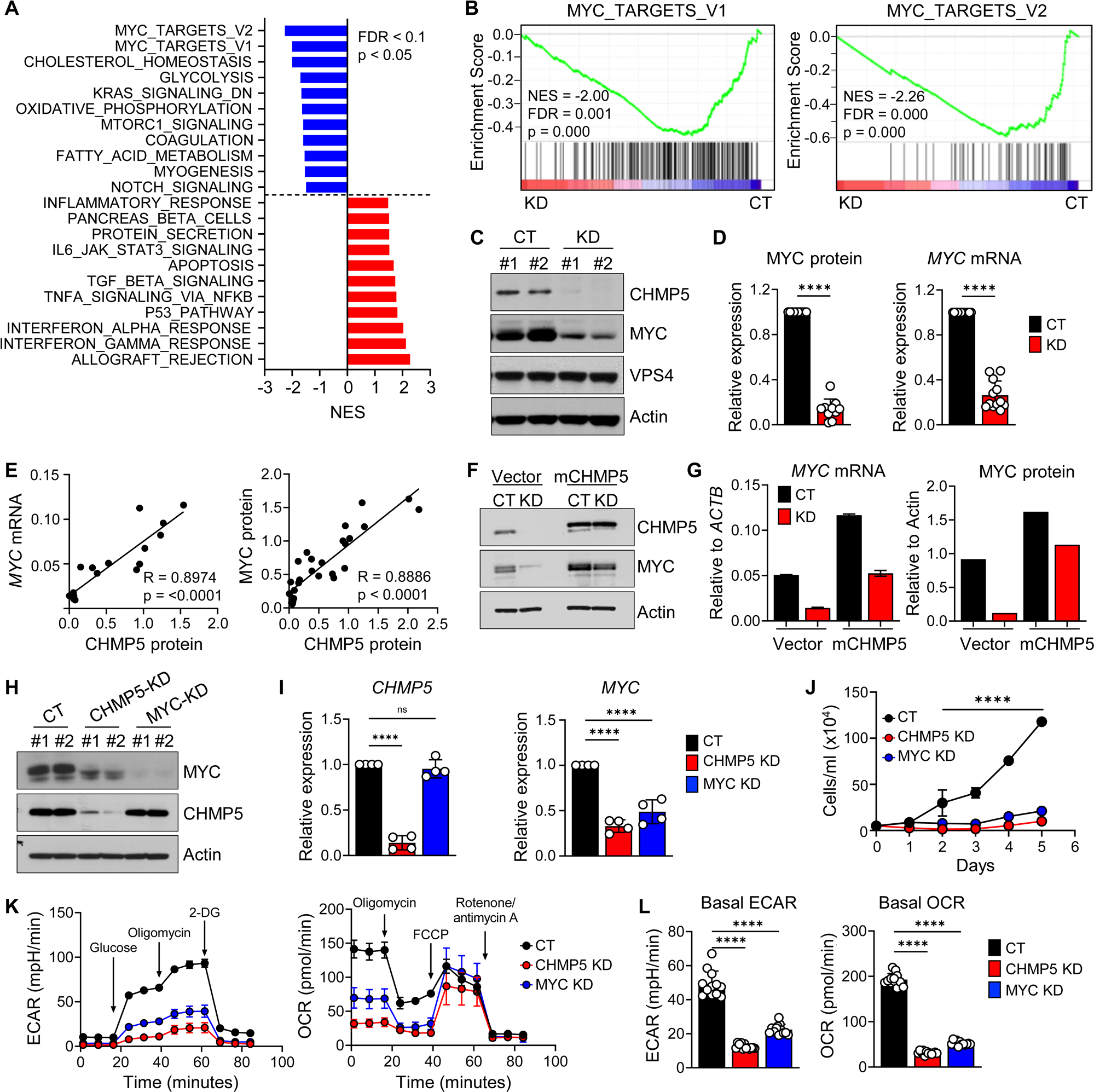
CHMP5 promotes a T-ALL transcriptional program exemplified by MYC. (A) Hallmark pathway enrichment scores of DEGs (fold-change ≥ 1.2 ; adjusted p < 0.05) between control (CT) versus CHMP5-deficient (KD) CUTLL1 cells after 5 days of selection with puromycin. (B) Hallmark GSEA plots of MYC-target gene pathways in CT versus KD CUTLL1 cells. (C) Western blot of the indicated proteins in CUTLL1 whole cell lysates. (D) MYC protein and mRNA expression relative to Actin and normalized to CT CUTLL1 cells. Data are presented as average (± SD) of replicates pooled from 5 independent experiments. Student’s t-test: ****, p < 0.0001. (E) Correlation between CHMP5 protein and *MYC* mRNA (left) and MYC protein (right). Data points are averaged replicates pooled from 5 independent experiments. Rho (R) and p-values determined by Pearson Correlation test. (F) Western blot of CT and KD CUTLL1 cells transduced with empty vector (Vector) or murine CHMP5 lentivirus (mCHMP5). (G) Quantification of MYC protein and mRNA relative to Actin of cells from (F). Data are representative of two experiments. (H) Western blot of CUTLL1 cells transduced two independent shRNAs for control (CT), CHMP5, or MYC. (I) mRNA expression of *CHMP5* and *MYC* relative to *ACTB* and normalized to CT. Data are presented as average (± SD) of biological replicates pooled from two independent experiments. One-way ANOVA: ****, p < 0.0001; ns, not significant (p > 0.05). (J) CUTLL1 growth kinetics determined by trypan blue counting. Data are average (± SD) cell numbers in triplicate wells. (K) Extracellular acidification rate (ECAR, left), and oxygen consumption rate (OCR, right) kinetics. Data are presented as mean (± SD) of 6 replicates of 2 independent shRNAs per group. 2-DG, 2-deoxyglucose; FCCP, carbonyl cyanide-4-(trifluoromethoxy) phenylhydrazone. (L) Basal ECAR and OCR values of indicated CUTLL1 cells. One-way ANOVA: ****, p < 0.0001.

Using two independent *CHMP5* targeting shRNAs we confirmed that *MYC* transcripts and MYC protein were indeed drastically decreased in CHMP5-deficient T-ALL cells (**Figures 1C** and **1D**). Control and CHMP5-deficient T-ALL cells showed comparable MYC protein stability in cycloheximide (CHX) chase assays and proteasome inhibition by MG132 failed to restore its MYC levels (**Figures S1E** to **S1G**), implying that CHMP5 likely controlled MYC expression at the level of transcription, in contrast to its post-translational activity in thymocytes and osteoclasts^29,30^. Of note, *MYC* transcripts and MYC protein levels positively correlated with the amount of CHMP5 proteins present in T-ALL cells (**Figure 1E**), suggesting a quantitative effect of CHMP5 on MYC expression in these human T-ALL cells.

To validate the specificity of CHMP5 loss on MYC expression, we transduced our CHMP5-deficient human T-ALL cells with control (“Vector”) or lentiviruses encoding murine *Chmp5* (mCHMP5) which is 99% identical in amino acid sequence to human CHMP5^33^. Murine CHMP5 not only restored *MYC* mRNA and MYC protein expression (**Figures 1F** and **1G**) but also rescued downstream MYC-dependent pathways like mitochondria oxidation and endoplasmic reticulum homeostasis in CHMP5-KD cells (**Figure S1H**). Interestingly, CHMP5 overexpression (due to transduced mCHMP5) further increased MYC protein levels in CHMP5-sufficient (CT) T-ALL cells (**Figures 1F** and **1G**), corroborating that CHMP5 quantitatively promoted MYC expression. Together, these findings uncover a previously unappreciated requirement for the ESCRT protein CHMP5 in promoting the ICN1-driven gene program exemplified by MYC in human T-ALL cells.

### CHMP5 deficiency phenocopies MYC deficiency in T-ALL cells

The proto-oncogene MYC is a master transcription factor of multiple genes involved in metabolism, protein synthesis and proliferation and is dysregulated in many cancers^34^, including T-ALL^5,17,35^. As “MYC targets” were the topmost downregulated pathways in CHMP5-deficient human T-ALL cells, we sought to further clarify the relationship between CHMP5 deficiency and MYC deficiency in these cells. To this end, we compared the effect of shRNA-mediated CHMP5 (CHMP5-KD) and MYC (MYC-KD) depletion on CUTLL1 T-ALL cells. Whereas loss of CHMP5 caused MYC downregulation, MYC depletion had no impact on CHMP5 expression (**Figures 1H** and **1I**), establishing CHMP5 function upstream of the MYC pathway in these T-ALL cells.

Consistent with a role of MYC in promoting cell cycle progression and metabolism^34^, *in vitro* growth kinetics of both MYC-KD and CHMP5-KD cells were equally blunted, accompanied by markedly reduced expression of glycolytic and oxidative phosphorylation genes like *PHB1, C1QBP, LDHA,* and *HK2* (**Figures 1J** and **S1I**). Correspondingly, CHMP5 depletion, like MYC deficiency, impaired energy metabolism as T-ALL cells lacking either protein displayed reduced basal and induced glycolytic capacity (ECAR) and mitochondrial respiration (OCR) (**Figures 1K** and **1L**). Further, both CHMP5-and MYC-deficient T-ALL cells displayed comparably diminished ER biogenesis, mitochondria reactive oxygen species (surrogate for oxidative phosphorylation), cell size, and protein synthesis (**Figures S1J**), cellular processes dictated by MYC^34^. Together, these data support MYC deficiency as a phenotype of CHMP5-deficient T-ALL cells and that CHMP5 promotes a transcriptional machinery driving MYC expression in these T-ALL cells.

### Identification of a nuclear fraction of CHMP5

The majority of ESCRT proteins, including CHMP5, are thought to be cytoplasmic^24,25,27^. However, the massive changes in the transcriptome brought about by CHMP5-deficiency raised the possibility that it might regulate gene transcription in the nucleus. Therefore, to understand how it influenced T-ALL gene transcription we first sought to clarify CHMP5’s cellular localization in T-ALL cells. Remarkably, we found that in addition to its previously reported cytosolic localization CHMP5 also localized to the nucleus in human T-ALL cell lines (CUTLL1, SUPT1) and primary patient-derived T-ALL cells (**Figure 2A**) and was also present in normal human T cells (**Figure S2A**). Overall, nuclear CHMP5 was ∼20-40% of cytosolic CHMP5 levels (**Figure 2A** and **S2A**).

**Figure 2.**
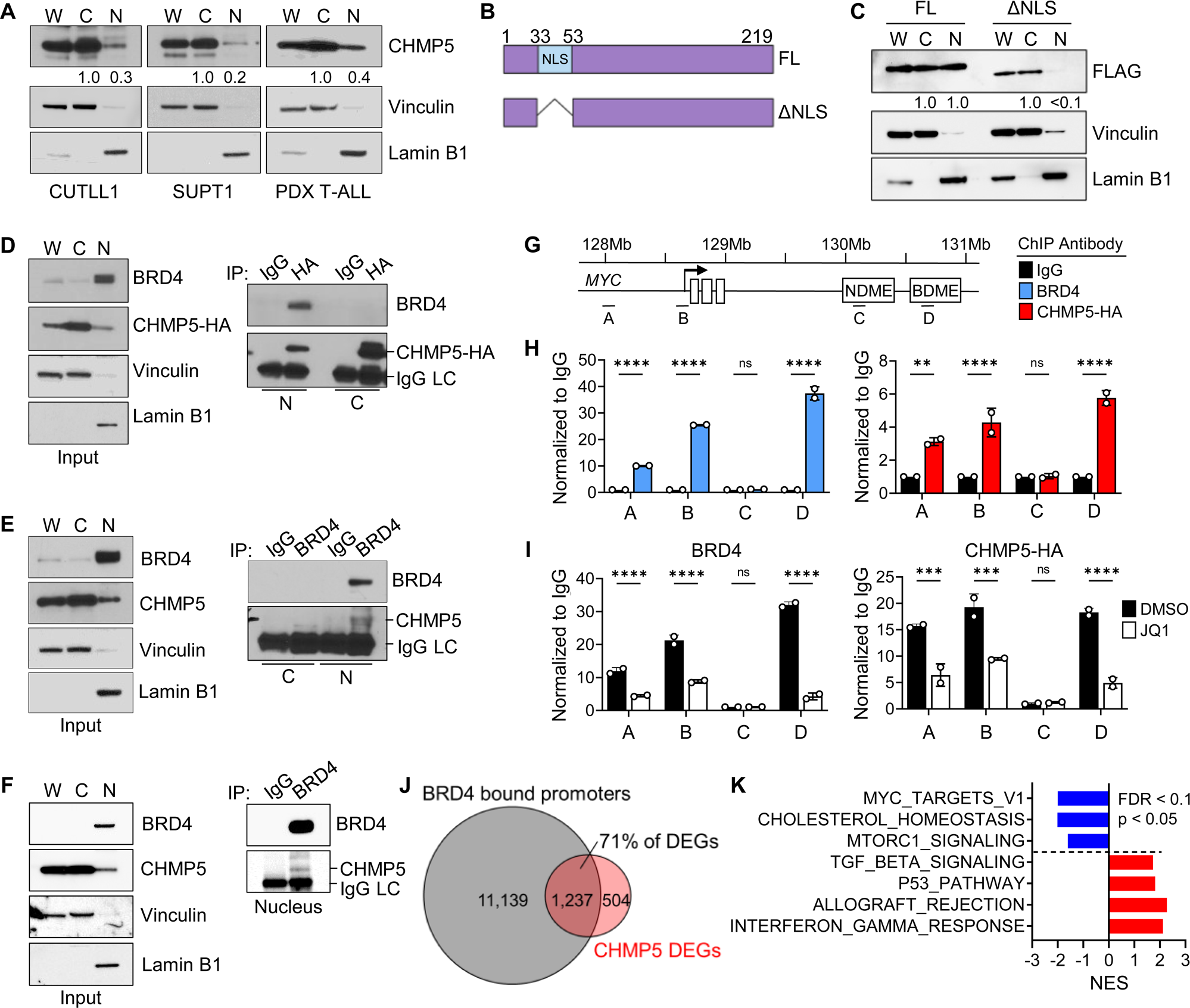
Identification of nuclear CHMP5-BRD4 interaction on chromatin. (A) Western blot of fractionated CUTLL1, SUPT1, and patient derived xenograft (PDX) T-ALL cells. W, whole cell; C, cytoplasmic; and N, nuclear lysates. CHMP5 band intensities relative to the cytoplasmic band are indicated. (B) Schematic representation of the NLS truncation mutant of human CHMP5. (C) Western blot of fractionated CUTLL1 cells transduced with FLAG-tagged full-length CHMP5 (FL) or ΔNLS mutant encoding lentivirus. FLAG (CHMP5) band intensities relative to the cytoplasmic band are indicated. (D) Western blot of fractionated CUTLL1 cells transduced with CHMP5-HA subjected to immunoprecipitation with isotype (IgG) or anti-HA antibodies. (E) Western blot of fractionated CUTLL1 cells subjected to immunoprecipitation with isotype (IgG) or anti-BRD4 antibodies. (F) Western blot of fractionated PDX T-ALL cells subjected to immunoprecipitation with isotype (IgG) or anti-BRD4 antibodies. (G) Schematic of the *MYC* gene locus indicating ChIP-qPCR primer binding: A, enhancer; B, promoter; C, NDME (NOTCH-dependent *MYC* enhancer); D, BDME (BRD4-dependent *MYC* enhancer). (H) ChIP-qPCR of anti-BRD4 and CHMP5-HA at the *MYC* locus normalized to isotype (IgG). Data points are technical replicates representative of 2 independent experiments. 2-way ANOVA: ****, p < 0.0001. (I) ChIP-qPCR of anti-BRD4 and CHMP5-HA normalized to IgG from CUTLL1 cells treated with vehicle (DMSO) or 500 nM of JQ1 for 18 hours. Data points are technical replicates representative of 2 independent experiments. 2-way ANOVA: ****, p < 0.0001. (J) Venn-diagram of BRD4 bound genes (determined by ChIP-seq, GSE51800) and DEGs from CT versus KD CUTLL1 cells. (K) Pathway analysis of the overlapping BRD4-bound genes and DEGs from CT versus KD CUTLL1 cells in (J).

To determine the basis of its nuclear localization, we examined the amino acid sequence of CHMP5 across various species for a nuclear localization signal (NLS). Notably, the N-terminus of CHMP5 in jawed vertebrates (including mice and humans), in which CHMP5 protein sequence is highly conserved, contained a putative bipartite NLS^36^ that was absent in invertebrate eukaryotes (**Figure S2B** and **S2C**). To verify this putative NLS we deleted the corresponding amino acid residues (**Figure 2B**) and assessed the localization of NLS-deficient (ΔNLS) CHMP5 proteins. Indeed, unlike full-length CHMP5 (FL), ΔNLS-CHMP5 largely failed to localize to the nucleus in CUTLL1 T-ALL cell (**Figure 2C**), indicating that, at least in T-ALL cells, CHMP5 proteins are imported into and localize to the nucleus.

### Nuclear CHMP5 interacts with BRD4

The unexpected nuclear localization of CHMP5 prompted us to hypothesize that CHMP5 directly mediated a mechanism that regulated transcription of T-ALL genes like MYC. Because in activated NOTCH1-induced T-ALL, ICN1 drives transcription in cooperation with BRD4 and MYC^17–19^, we investigated whether nuclear CHMP5 interacted with these transcription factors. Given the lack of suitable immunoprecipitation antibodies for CHMP5, we transduced CUTLL1 T-ALL cells with hemagglutinin (HA)-tagged CHMP5 and performed anti-HA immunoprecipitation from these cells. While CHMP5 did not interact with ICN1 or MYC (**Figure S2D**), we found that it associated with endogenous BRD4 in nuclear lysates in T-ALL cells (**Figure 2D**). We confirmed this CHMP5-BRD4 interaction in independent reverse immunoprecipitation of endogenous BRD4 in nuclear lysates from human T-ALL cell lines (CUTLL1 and Loucy), primary PDX human T-ALL (**Figures 2E, 2F** and **S2E**), and in HEK293T cells co-transfected with plasmids encoding these proteins (**Figure S2F**). Importantly, in a cell-free assay, recombinant BRD4 pull-downed CHMP5 (**Figure S2G**), indicating a direct interaction between both proteins.

To determine whether nuclear CHMP5 bound BRD4 on chromatin we performed anti-BRD4 and anti-HA (CHMP5) chromatin immunoprecipitation followed by qPCR (ChIP-qPCR) on T-ALL cells transduced with HA-tagged CHMP5. We confirmed that both antibodies significantly pulled-down chromatin relative to isotype antibody (**Figure S2H**), validating our anti-HA ChIP approach. Focusing on the *MYC* gene locus, ChIP-qPCR revealed CHMP5 binding on chromatin at sites that overlapped with BRD4 binding, including at the *MYC* enhancer, promoter, and a BRD4-dependent super-enhancer (“BDME”)^21^ (**Figures 2G** and **2H**). CHMP5 binding was indistinguishable from isotype (IgG) antibody at the ICN1-specific (“NDME”) super-enhancer which is not bound by BRD4^21^ (**Figures 2G** and **2H**), suggesting that CHMP5 bound chromatin at the *MYC* gene through its association with BRD4. Accordingly, treatment with the BET inhibitor JQ1 which releases BRD4 from chromatin^37^ significantly reduced CHMP5 binding across the *MYC* locus (**Figure 2I**).

That CHMP5 associated with BRD4 on chromatin regulatory elements at the *MYC* locus raised the possibility that it regulated transcription in T-ALL cells through a BRD4-dependent mechanism. In line with this, we found that the majority (∼71%) of DEGs in CHMP5-deficient T-CUTLL1 T-ALL cells have BRD4 bound promoters (GSE51800)^38^ and were genes that mediated the topmost perturbed pathways in these CHMP5-deficient T-ALL cells (**Figures 2J** and **2K**).

### CHMP5 promotes Pol II pause-release at BRD4-dependent T-ALL genes

BRD4 has been shown to promote transcriptional elongation by facilitating promoter-proximal Pol II pause release^8,11,39,40^. Given the interaction between BRD4 and CHMP5, and that a substantial number of DEGs in CHMP5-deficient T-ALL cells are BRD4-regulated genes (**Figure 2J**), we asked whether CHMP5 promoted Pol II pause release. To this end, we performed genome-wide ChIP-seq for Pol II and BRD4 in CHMP5-depleted (KD) or control (CT) CUTLL1 T-ALL cells. ChIP-seq analysis showed a modest increase in BRD4 binding at transcriptional start sites (TSS) but largely unchanged binding across, gene bodies, and transcriptional end sites (TES) in CHMP5-depleted T-ALL cells (**Figures S3A** and **S3B**). However, in parallel, CHMP5-depleted T-ALL cells displayed decreased Pol II occupancy at the TES with a corresponding increase of Pol II binding at promoters (**Figures 3A** to **3C** and **S3C**), indicating impaired Pol II pause-release. Importantly, the loss of Pol II at the TES correlated with gene expression, as reduction in Pol II density at the TES was significantly higher in downregulated DEGs compared to upregulated DEGs in KD T-ALL cells (**Figure S3D**). In light of this finding, as BRD4 travels with the Pol II during transcription elongation^40–42^, increased BRD4 occupancy at promoters may reflect its binding to stalled Pol II.

**Figure 3.**
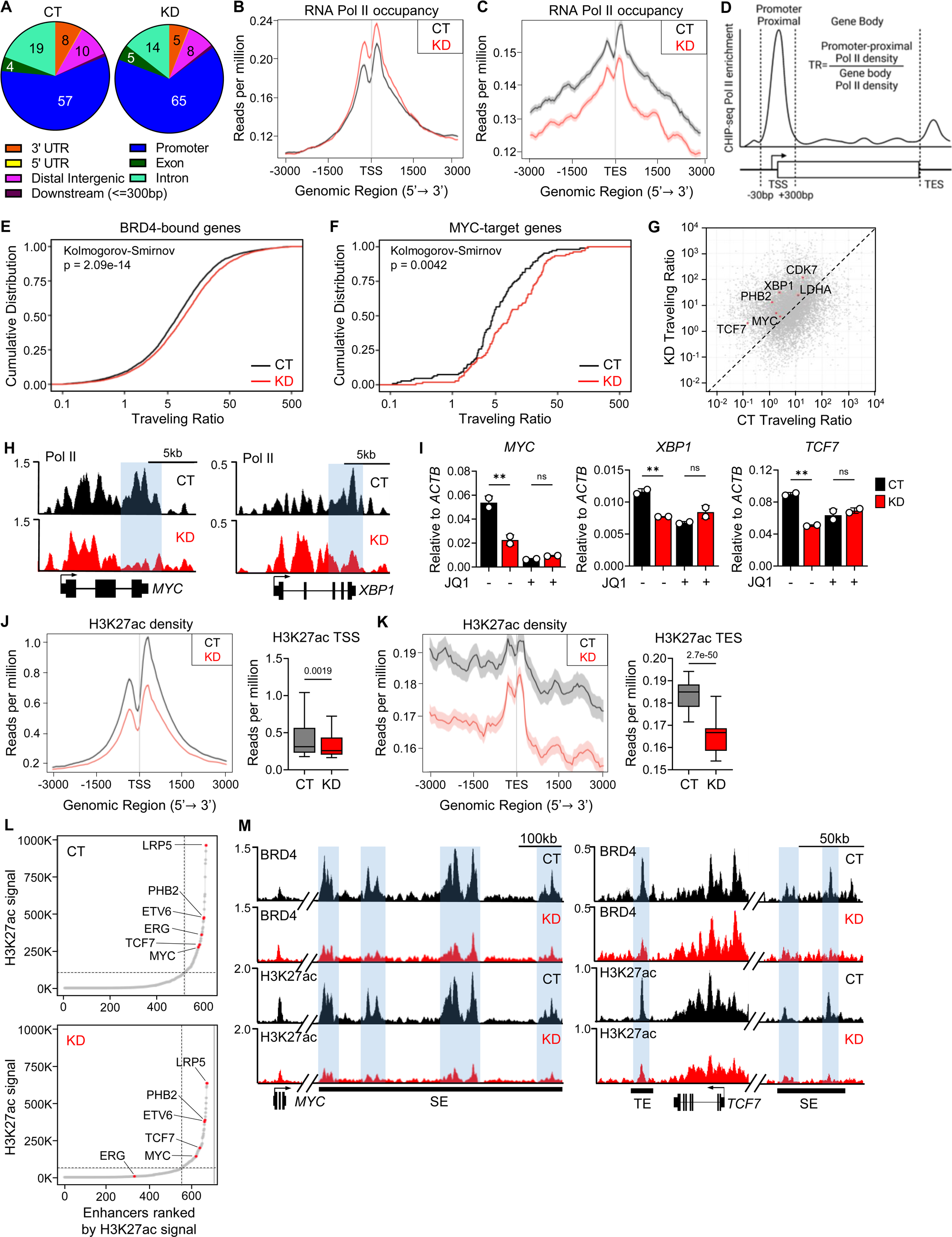
CHMP5 mediates BRD4-driven Pol II pause release and super enhancer formation. (A) Pie chart of genome-wide Pol II binding in control (CT) and CHMP5-depleted (KD) CUTLL1 cells determined by ChIP-seq. (B) Metaplot of Pol II density at TSS of all genes from (A). p-value, Student’s t-test. (C) Metaplot of Pol II density at TES of all genes from (A). p-value, Student’s t-test. (D) Schematic for the calculation of Pol II traveling ratio. (E-F) Pol II traveling ratio of CT and KD CUTLL1 cells for BRD4-bound genes (E) and MYC-Target genes (F). (G) Dot plot comparison of Pol II traveling ratios between CT and KD cells highlighting BRD4 and MYC-target genes. (H) Pol II binding tracks in CT and KD CUTLL1 cells at the *MYC* and *XBP1* gene loci. 3’-end of genes are highlighted. (I) mRNA expression of *MYC, XBP1,* and *TCF7* in CUTLL1 CT and KD cells treated with DMSO (−) or JQ1 (+) for 18 hours. Data points are technical replicates. Student’s t-test: **, p < 0.01; ns, not significant. (J) Metaplot and box plot of H3K27ac density at TSS of active genes determined by ChIP-seq. p-value, Student’s t-test. (K) Metaplot and box plot of H3K27ac density at TES of active genes determined by ChIP-seq. p-value, Student’s t-test. (L) Hockey stick plots of ranked genome-wide H3K27ac signals in CT (top) and KD (bottom) CUTLL1 cells. Positions of key T-ALL genes are highlighted. (M) BRD4 and H3K27ac ChIP-seq tracks at the *MYC* and *TCF7* gene loci in CUTLL1 cells. SE, super-enhancer; TE, typical enhancer.

To quantify the extent to which CHMP5 loss impacted Pol II pause release, we calculated genome-wide Pol II traveling ratios (TR), the relative ratio of Pol II density at promoter-proximal regions relative to its density across the gene body (**Figure 3D**)^8,43^. Overall, we found higher TRs for BRD4-target genes^38^ in CHMP5-deficient T-ALL cells (**Figure 3E**) confirming impaired Pol II pause release at promoters and decreased transcriptional elongation of these T-ALL genes. Since BRD4 also transactivates T-ALL genes in cooperation with MYC, another Pol II pause release factor^44^, we also determined that the TRs for MYC target genes were higher in CHMP5-deficient T-ALL cells (**Figure 3F**). Correlation of the TR of all genes in control (CT) versus CHMP5-deficient (KD) cells highlighted an increased TR of key T-ALL genes including essential pro-leukemogenic genes like *MYC*, *TCF7, PHB2*, *CDK7*, and *XBP1* (**Figure 3G**). Accordingly, Pol II density at the 3’-ends of these genes (exemplified by *MYC* and *XBP1*) was markedly diminished (**Figure 3H**).

Reflecting their BRD4-dependency, mRNA levels for T-ALL genes that displayed Pol II stalling (e.g., *MYC*, *XBP1* and *TCF7*) were comparably downregulated by loss of CHMP5 and by the BET inhibitor JQ1 (**Figure 3I**). In fact, despite intact (or increased) BRD4 binding at promoter-proximal and gene body in these cells (**Figures S3A** and **S3B**), JQ1 treatment had no additional effect on downregulating the BRD4-target genes *XBP1* and *TCF7* in CHMP5-deficient T-ALL cells (**Figure 3I**). Altogether, these findings are consistent with a model in which CHMP5 promotes Pol II pause release and transcription of BRD4-dependent genes in NOTCH1-driven T-ALL.

### Loss of CHMP5 impairs H3K27 acetylation and disrupts super-enhancers

Depletion of CHMP5 did not impact promoter-proximal (TSS) BRD4 binding, which remained largely unchanged in CHMP5-KD T-ALL cells (**Figures S3A** and **S3B**). Therefore, to gain insight into how CHMP5 promoted transcription of BRD4-dependent T-ALL genes like MYC, we wondered if it instead controlled BRD4 binding at distal enhancers defined by hyperacetylated histone modifications, including H3 lysine 27 acetylation (H3K27ac)^45,46^. Notably, recruitment of BRD4 to these hyperacetylated enhancers and super-enhancers promote Pol II pause release and transcriptional elongation^8,40,47,48^. We performed H3K27ac specific ChIP-seq and found diminished H3K27ac density at the TSS, gene bodies, and TES of active genes in CHMP5-KD T-ALL cells (**Figures 3J, 3K** and **S3E**). We first confirmed that actively transcribed genes (identified by positive H3K27ac signal), showed impaired Pol II traveling (**Figure S3F).**

Following previously reported methods^45,46^, we then defined enhancers and super-enhancers by ranking H3K27ac signal intensity across the genome. As reported in cancer and stem cells^45,46^, “hockey stick” plots showed that H3K27ac signals were asymmetrically distributed across the genome, with disproportionately higher density at super-enhancers of T-ALL genes like *MYC*, *TCF, ERG,* and *ETV6*^40^ (**Figure 3L**). Importantly, consistent with their downregulated expression (**Figure S3G**), we found drastically diminished H3K27ac density across these key T-ALL genes, with significant loss of H3K27ac and BRD4 at both typical enhancers and super-enhancers in CHMP5-deficient T-ALL cells (**Figures 3M** and **S3H**). Thus, CHMP5 promotes H3K27ac that defines enhancer and super-enhancers to which BRD4 binds to facilitate Pol II pause release and transcription^8,40,47,48^. Of note, despite reduced H3K27ac density at TSSs, CHMP5 deficiency did not reduce BRD4 binding at TSS, likely reflecting BRD4 binding to promoter-bound proteins like Pol II. Furthermore, that CHMP5 deficiency largely only impaired BRD4 binding at enhancers and super-enhancers, implied that it functioned selectively to regulate epigenetic events at distal regulatory elements and not promoters.

### Potentiation of BRD4-p300 interaction by CHMP5

The histone acetyl transferase (HAT) p300 (with its structural homolog CBP) is a major catalyst of H3K27ac of enhancers and super-enhancers^49–51^. Studies in normal and cancer cells suggest a model in which p300 recruits BRD4 to hyperacetylated enhancers and super-enhancers and in turn, BRD4 augmented p300’s HAT activity, a positive feed-forward mechanism that enriches hyperacetylation of enhancers and super-enhancers^52–54^. We thus asked if CHMP5 regulated the BRD4-p300 axis to promote H3K27ac and assessed the impact of CHMP5 deficiency on BRD4-p300 interaction. While BRD4’s interaction with components of the transcriptional machinery (e.g., MED1, Pol II) were intact, its interaction with p300 was significantly reduced in CHMP5-deficient T-ALL cells (**Figure 4A** and **4B**). This was despite an overall increase in total p300 expression in CHMP5-deficient cells (**Figures 4A** and **S4A**).

**Figure 4.**
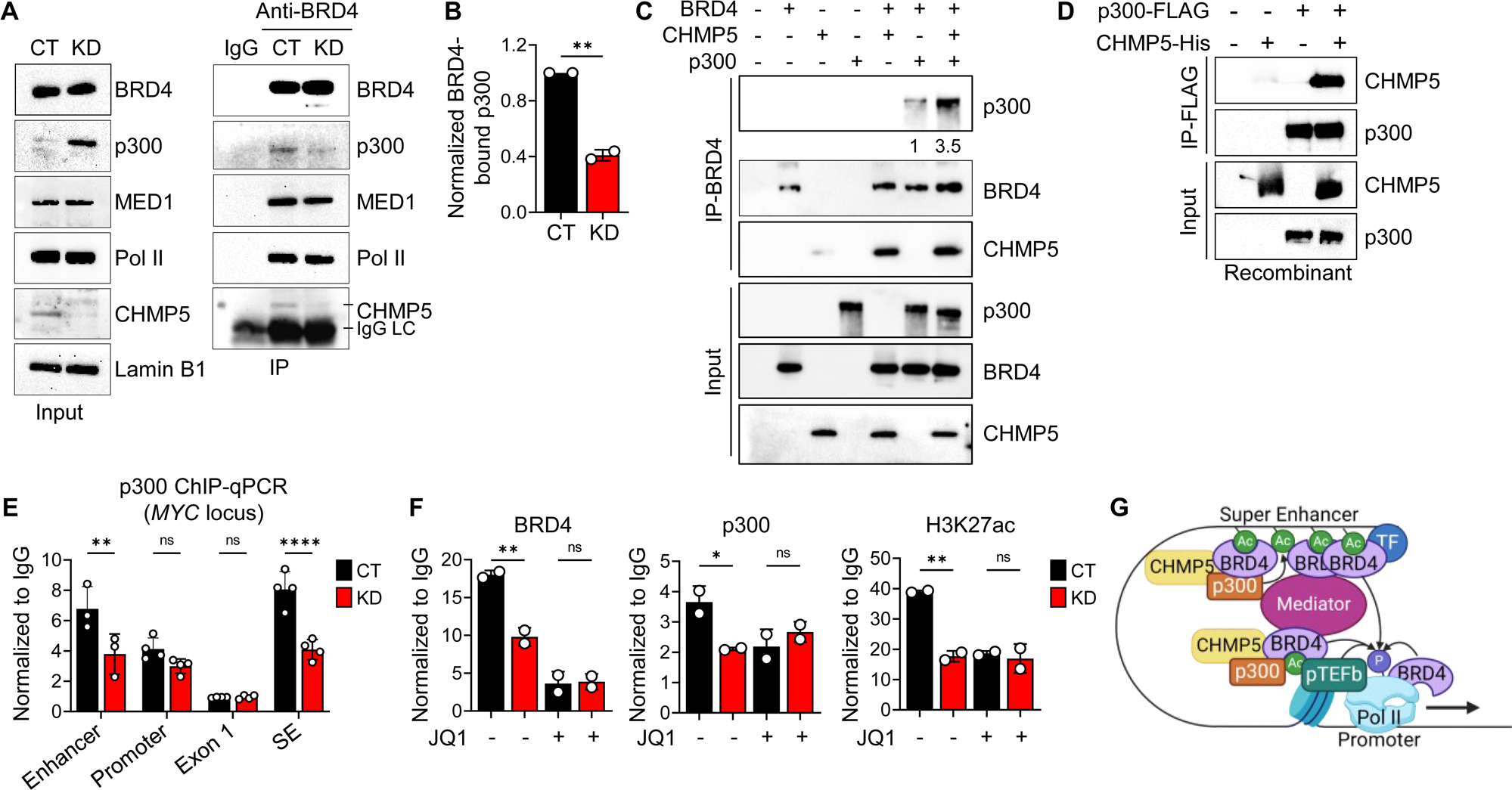
CHMP5 promotes the interaction between BRD4 and p300. (A) Immunoprecipitation with isotype (IgG) or anti-BRD4 antibody from CT and KD CUTLL1 nuclear lysate. (B) Quantification of p300 bound to BRD4 from (**A**) calculated as p300-to-BRD4 ratio and normalized to control (CT) cells. Data are average (± SD) of two independent experiments. Student’s t-test: **, p < 0.01. (C) Recombinant BRD4, p300, and CHMP5 immunoprecipitation with anti-BRD4 antibody. Ratio of p300 bound to BRD4 (quantified by densitometry) relative to no-CHMP5 lane 6 is indicated. Data is representative of 2 independent experiments. (D) Recombinant p300-FLAG and CHMP5-His immunoprecipitation with anti-FLAG antibody. (E) ChIP-qPCR of p300 at the *MYC* locus normalized to isotype (IgG) in CT and KD CUTLL1 cells. Data are average (± SD) of technical replicates from two independent experiments. 2-way ANOVA: **, p < 0.01; ****, p < 0.0001; ns, p > 0.05. SE, super-enhancer. (F) ChIP-qPCR of BRD4, p300, and H3K27ac at the *MYC* SE in CUTLL1 cells treated with DMSO (−) or 500 nM of JQ1 (+) for 18 hours. Data are average (± SD) of technical replicates. One-way ANOVA: *, p < 0.05; **, p < 0.01; ns, p > 0.05. (G) Schematic depiction of how the CHMP5-mediated p300-BRD4 interaction promotes acetylation and productive transcription.

To directly assess the impact of CHMP5 on the BRD4-p300 interaction, we quantified recombinant BRD4 and p300 binding in the presence and absence of recombinant CHMP5 protein. and found that more p300 proteins co-immunoprecipitated with BRD4 when CHMP5 was present (**Figure 4C**). Immunoprecipitation studies in HA-CHMP5-expressing T-ALL cells showed p300 association with CHMP5 in nuclear lysates from CUTLL1 T-ALL cells (**Figure S4B**) and in cell-free assays, recombinant CHMP5 immunoprecipitated with p300 (**Figure 4D**), consistent with a direct binding between both proteins. As it also directly binds BRD4 (**Figure S2G**), these findings suggest that CHMP5 promoted H3K27 acetylation at regulatory enhancers and super-enhancers by augmenting the BRD4-p300 interaction, an interaction that potentiates both p300 and BRD4-mediated H3K27ac at enhancers and super-enhancers^53^.

Consistent with a model in which CHMP5 selectively regulated epigenetic events at distal regulatory elements, loss of CHMP5 resulted in reduced p300 binding at the *MYC* enhancer and super-enhancer (BDME), while p300 binding remained unchanged at the promoter (**Figure 4E**). Of note, depletion of CHMP5 reduced p300 and H3K27ac at the *MYC* super-enhancer (BDME) to the same extent as treatment with the BET inhibitor JQ1 (**Figure 4F**), establishing CHMP5 upstream of the p300-BRD4 recruitment. Moreover, while JQ1 further decreased super-enhancer-bound BRD4 in CHMP5-deficient T-ALL cells, it had no additional impact on p300 and H3K27ac signals at the super-enhancer in CHMP5-deficient T-ALL cells (**Figure 4F**). These findings suggest that CHMP5 mediates the H3K27ac effected by the p300-BRD4 axis at super-enhancers.

Based on findings from our ChIP-seq and protein interaction studies, we envision a model (**Figures 4G** and **S4C**) in which nuclear CHMP5 functions as a positive regulator of the sequential events that establish distal chromatin modifications, including acetylation of enhancers and super-enhancers, which are bound by BRD4 and p300 to facilitate Pol II pause release and transcriptional elongation^51,55^. Specifically, in CHMP5-sufficient T-ALL cells, p300-catalyzed H3K27ac recruits initial BRD4 binding via its bromodomain and in turn, chromatin-bound BRD4 recruits additional p300 proteins, which catalyzes further H3K27ac reactions and enrichment at enhancers and super enhancers. Interaction of factors at such distal regulatory elements and promoter-proximal factors orchestrate productive transcriptional elongation (**Figure S4C**). Conversely, in CHMP5-deficient T-ALL cells which have diminished p300-BRD4 interaction and recruitment to chromatin, enhancers and super-enhancers are hypoacetylated and transcriptional elongation is impaired (**Figure S4C**).

### High CHMP5 expression is prognostic in human T-ALL

Our findings that CHMP5 mediated the p300-BRD4 interaction that enabled expression of key T-ALL genes such as MYC suggest a significant role for CHMP5 in T-ALL pathogenesis. Thus, we wished to understand the significance of CHMP5 expression in primary human T-ALL disease. Interestingly, across hematological and solid cancers, lymphoid leukemias expressed the highest amount of CHMP5 proteins (**Figure S5A**). Specific comparison of CHMP5 expression in pediatric T-ALL samples versus normal thymocytes (GSE33470, GSE33469)^56,57^ revealed higher *CHMP5* transcripts in T-ALL cells (**Figure 5A**) which translated to >5-fold more CHMP5 proteins in primary human T-ALL relative to healthy T-cells (**Figures 5B** and **5C**; **Table 1**). Human T-ALL cell lines similarly expressed more CHMP5 protein relative to healthy T-cells (**Figures S5B** and **S5C**). Of note, consistent with a unique role for CHMP5 in T-ALL, expression of other ESCRT proteins like CHMP1A (which can also localize to the nucleus^58^) and VPS4 were comparable in normal T-cells and T-ALL cells (**Figures 5B, S5B** and **S5C**).

**Figure 5.**
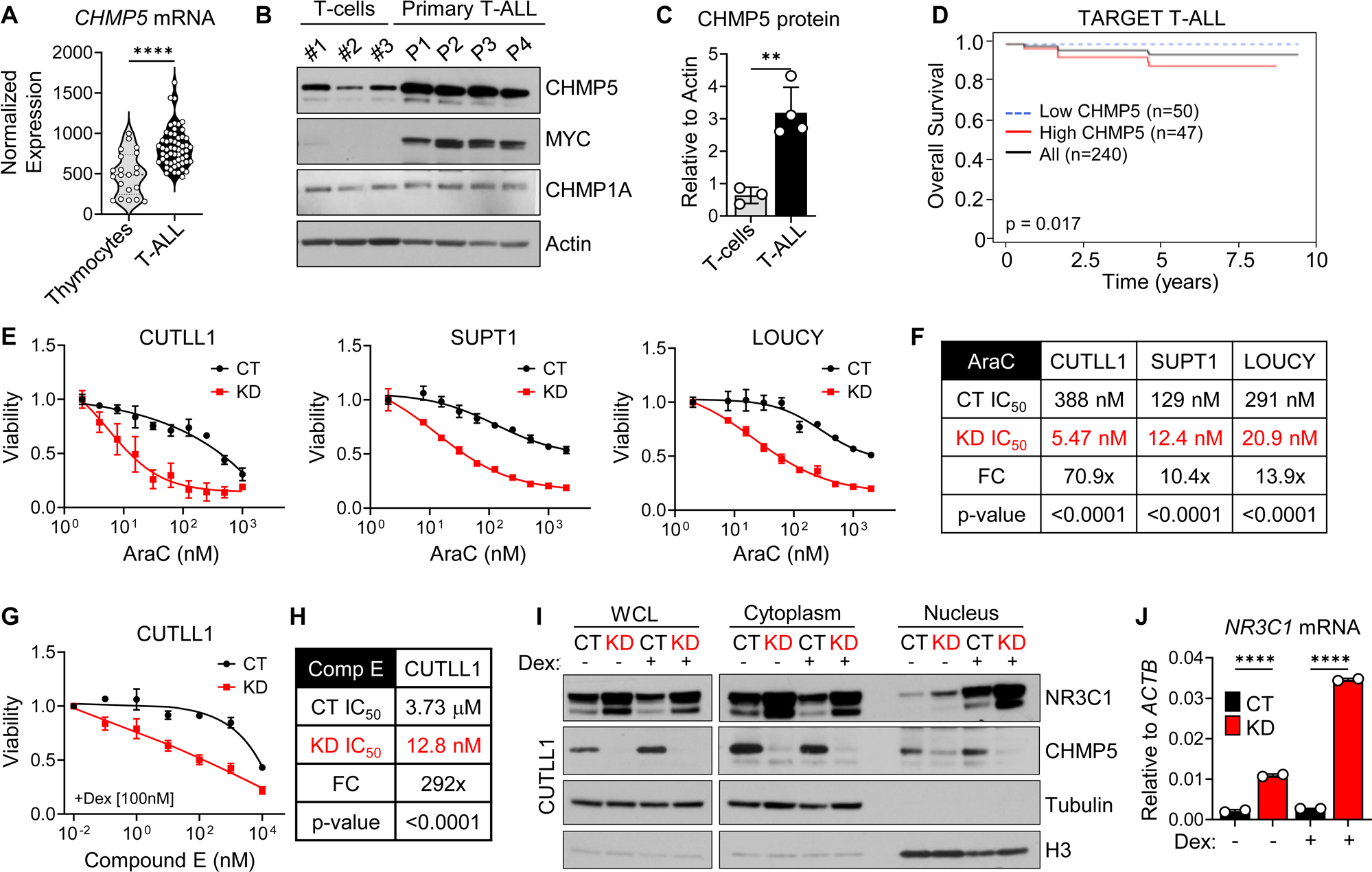
Significance of CHMP5 expression in human T-ALL disease prognosis and chemoresistance. (A) *CHMP5* mRNA expression in normal thymocytes (n = 21) and primary T-ALL (n = 57) samples (GSE33470, GSE33469). Student’s t-test: ****, p < 0.0001. (B) Western blot of indicated proteins in T cells from healthy donors (n = 3) and primary human T-ALL samples (n = 4). (C) Quantification of CHMP5 protein relative to Actin from (B). Student’s t-test: **, p < 0.01. (D) Overall survival of pediatric T-ALL patients (TARGET T-ALL) expressing high (top 20%) and low (bottom 20%) levels of CHMP5. p = 0.017, Log-rank test. (E) Viability of control (CT) and CHMP5-depleted (KD) CUTLL1 (left), SUPT1 (middle), LOUCY (right) cells treated with cytarabine (AraC) for 3 days. Data are presented as mean (± SD) of 3 replicates. Representative of 2 independent experiments. (F) IC_50_ for AraC in CUTLL1, SUPT1, and LOUCY cells from (E). IC_50_ calculated by non-linear best-fit analysis. p-value calculated by 2-way ANOVA. FC, fold-change in CT versus KD IC_50_. (G) Viability of CT and KD CUTLL1 cells treated with Compound E (Comp E) plus 100 nM dexamethasone (Dex) for 3 days. Data are presented as mean (± SD) of 3 replicates. (H) IC_50_ for Compound E in CT and KD CUTLL1 cells. IC_50_ calculated by non-linear best-fit analysis. p-value calculated by 2-way ANOVA. FC, fold-change in CT versus KD IC_50_. (I) Western blot of fractionated CT and KD CUTLL1 cells treated with vehicle (DMSO) or 1 μM dexamethasone (Dex) for 18 hours. (J) Expression of *NR3C1* in CUTLL1 cells from (I). Data are mean (± SD) of technical replicates. One-way ANOVA: ****, p < 0.0001.

**Table 1.** Patient information for primary T-ALL samples.

| Sample | Sex | Status at collection | Treatment | Active NOTCH1 | Phenotypic markers |
| --- | --- | --- | --- | --- | --- |
| P1 | male | Relapsed | Hydrea, leukopheresis | Yes | CD38+, CD34+ subset, CD3 int, CD5 dim, HLA-DR+ subset |
| P2 | male | De Novo | Hydrea, rasburicase | Yes | CD38+. CD34-, CD3+, CD5+, HLA-DR int |
| P3 | male | De Novo | Hydrea | Yes | CD38+, CD34+, CD3+, HLA-DR dim |
| P4 | male | De Novo | Allopurinol, leukopheresis | Yes | CD38+, CD34-, CD3-, CD5-, HLA-DR- |

To understand the implications of CHMP5 expression on human T-ALL disease, we evaluated the prognostic significance of *CHMP5* expression levels in the pediatric TARGET T-ALL dataset (dbGaP phs000464)^59^. While overall pediatric T-ALL patient survival was high in this cohort, we found that patients with the highest (top 20%) *CHMP5* expression had poorer overall survival (Log-rank test, *P* = 0.017) compared to patients with the lowest (bottom 20%) *CHMP5* expression in this cohort (**Figure 5D**). In line with their comparable expression in normal T-cells and T-ALL cells (**Figures 5B, S5B** and **S5C**), expression of VPS4A and CHMP1A did not correlate with T-ALL patient survival (**Figures S5D** and **S5E**), supporting the unique role for CHMP5 in T-ALL pathogenesis. Furthermore, *CHMP5* expression was significantly higher in T-ALL cells from adult patients that did not achieve complete remission^60^ (**Figure S5F**).

Interestingly, although our discovery of the critical role for CHMP5 in promoting T-ALL gene transcription were derived in the NOTCH1-driven human T-ALL model (CUTLL1), CHMP5 expression levels were prognostic in the TARGET T-ALL cohort that comprised patients with T-ALL disease with underlying mutations besides NOTCH1^59^. Moreover, high CHMP5 expression marked both NOTCH1 and non-NOTCH1-driven human T-ALL cell lines (**Figure S5B**). Together these observations reinforce that CHMP5 promoted expression of the T-ALL gene program by regulating a fundamental transcriptional mechanism -namely, the p300-BRD4 crosstalk-that is probably downstream of other T-ALL oncogenes besides ICN1 and shared across T-ALL subtypes.

### CHMP5 promotes chemoresistance in T-ALL

The poorer overall survival of T-ALL patients with the highest expression of CHMP5 (**Figure 5D**) suggested that CHMP5-regulated mechanisms dictate patient response to chemotherapy. To address this possibility, we assessed the impact of CHMP5 deficiency on T-ALL cell sensitivity to killing by chemotherapy and measured the viability of control (CT) and CHMP5-deficient (KD) human T-ALL cells after cytarabine (AraC) treatment *in vitro*. CHMP5-deficient CUTLL1 T-ALL cells showed significantly increased sensitivity with >10-fold reduction in IC_50_ for AraC (**Figures 5E** and **5F**). Reflecting that the CHMP5-BRD4-p300 axis is a general requirement in T-ALL cells, loss of CHMP5 also synergized with AraC in both activated NOTCH1-driven (CUTLL1 and SUPT1) and NOTCH1-independent (LOUCY) T-ALL cells (**Figures 5E** and **5F**).

NOTCH1-driven T-ALL cells like the CUTLL1 human T-ALL cell line are highly resistant to single or combination treatment with γ-secretase inhibitors (GSI) and glucocorticoids including dexamethasone (Dex) as ICN1-induced program represses expression of the glucocorticoid receptor NR3C1^61,62^. Therefore, to delineate CHMP5’s contribution to oncogene-specific chemoresistance we assessed the impact of CHMP5 deficiency on CUTLL1 T-ALL cells response to GSI+Dex treatment. Compared to control cells (CT), CHMP5-deficient (KD) CUTLL1 cells were strongly inhibited by GSI+Dex treatment (**Figures 5G** and **5H**). Increased sensitivity to GSI+Dex treatment was associated with higher expression and increased nuclear translocation of the glucocorticoid receptor NR3C1 (**Figures 5I)**, as well as increased expression of NR3C1-induced genes, including *NR3C1* and *BIM* (**Figures 5J** and **S5G)**.

Together, these drug response studies indicate that CHMP5 is part of the cellular mechanism that promotes chemoresistance in T-ALL cells. Interestingly, in contrast to its synergy with AraC and GSI+Dex that respectively target DNA replication and NOTCH1 processing pathways that are independent of the p300-BRD4 axis, CHMP5 deficiency did not synergize with the BET inhibitor JQ1 to kill T-ALL cells (**Figure S5H** and **S5I**). This is consistent with our model (**Figure S4C**) where CHMP5 functions upstream of and promotes p300 recruitment of BRD4 at enhancers and super-enhancers.

### CHMP5 is required for T-ALL initiation *in vivo*

To further understand the physiological significance of CHMP5 function in T-ALL disease, we investigated the contribution of CHMP5 to T-ALL development *in vivo*. We utilized a murine bone marrow chimera T-ALL model in which BM donor cells were transduced with retrovirus encoding oncogenic ICN1 that initiates a CD4^+^CD8^+^ subtype T-ALL disease^63,64^. Bone marrow (BM) cells (CD45.2^+^) enriched for c-Kit^+^ progenitors from littermate *Chmp5*^f/f^*Cd4*-Cre^−^ (WT) or *Chmp5*^f/f^ *Cd4*-Cre^+^ (KO) mice^30^ were transduced with bi-cistronic IRES retrovirus encoding ICN1, and a truncated nerve growth factor (NGFR) protein used to identify ICN1-transduced donor cells (**Figure S6A** and **S6B)**. Since Cre recombinase expression is driven by the *Cd4* promoter, *Chmp5* deletion is restricted to CD4-expressing cells^30^ in this murine T-ALL model. Transduction efficiency was confirmed by NGFR expression and transduced BM cells were transplanted into lethally irradiated congenic (CD45.1^+^) hosts along with congenic (CD45.1^+^) whole BM cells for hemogenic support (**Figure S6B** and **S6C**).

As expected^63,64^, chimera mice transplanted with ICN1-transduced WT BM cells (hereafter referred to as “WT” T-ALL mice) developed an aggressive, lethal (median survival, ∼5 weeks) and fully penetrant T-ALL disease characterized by marked leukocytosis, splenomegaly with noticeably disrupted white and red pulp zones, and tissue infiltration by leukemic cells (**Figures 6A** to **6C** and **S6D**). In striking contrast, chimera mice that were transplanted with ICN1-transduced KO donor BM cells (hereafter referred to a “KO” T-ALL mice) survived and did not develop any detectable T-ALL symptoms up to 12 weeks after transplantation (**Figures 6A** to **6C** and **S6D**).

**Figure 6.**
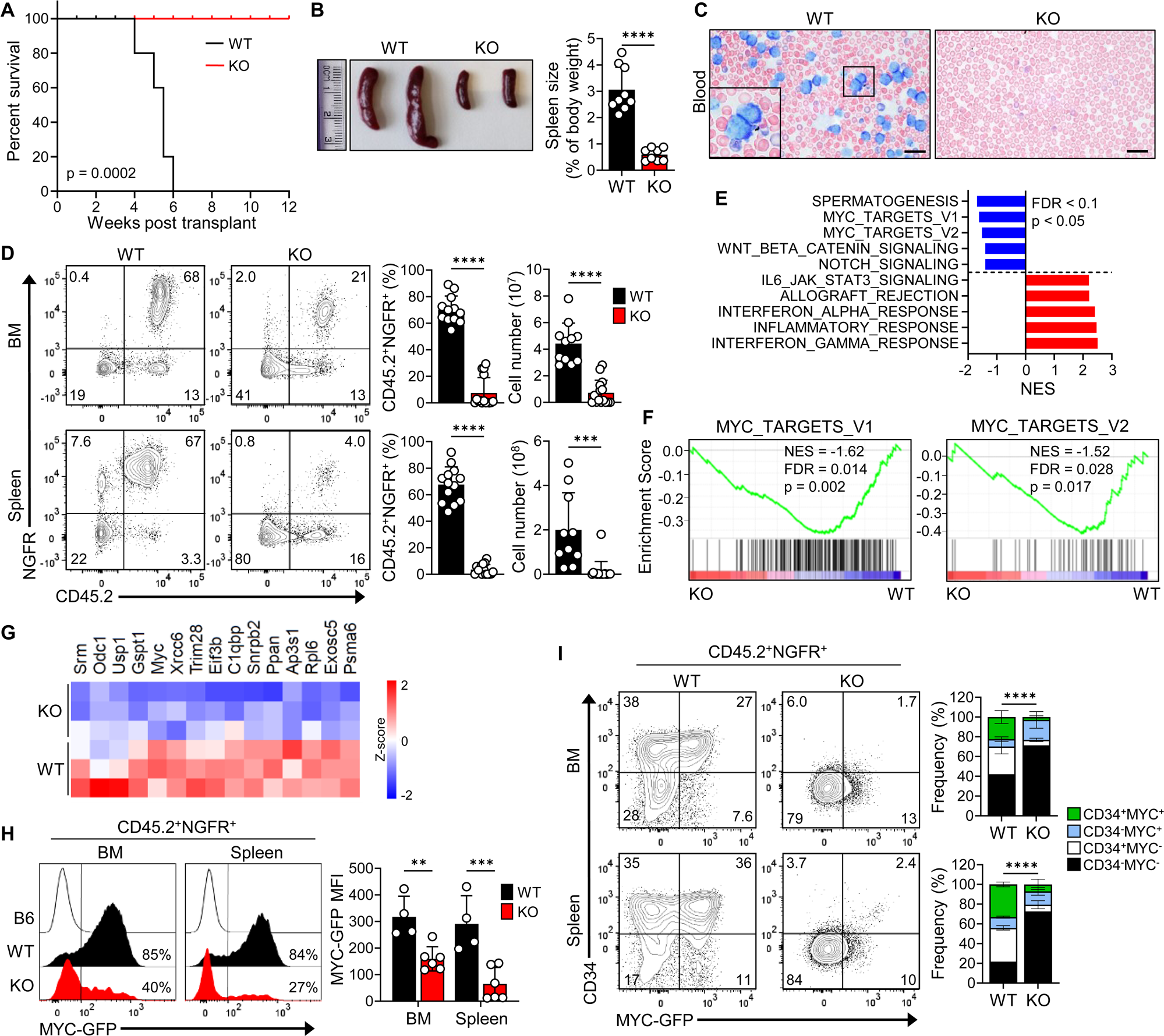
CHMP5 deficiency impairs T-ALL development and progression in vivo. (A) Kaplan-Meier survival curve of leukemia mice generated by injection of ICN1-NGFR-transduced *Chmp5*^f/f^*Cd4*-Cre^−^ (WT) or *Chmp5*^f/f^*Cd4*-Cre^+^ (KO) c-Kit-enriched bone marrow (BM cells). p, Mantel-Cox log-rank test; n = 9 mice/group. (B) Spleen images and average (± SD) spleen weights. Student’s t-test: ****, p < 0.0001; n = 9 mice/group, from 2 independent experiments. (C) Wright-Giemsa staining of blood from leukemia mice at 4 weeks post-transplant. Scale bar = 20 µm (D) CD45.2 and NGFR expression on BM (top) and spleen (bottom) cells at 4-weeks post-transplant with graphs of mean (± SD) frequency and numbers of CD45.2^+^NGFR^+^ cells. Student’s t-test: ***, p < 0.001; ****, p < 0.0001; n=11-15 mice/group, from 2 independent experiments. (E) Hallmark pathway enrichment scores of DEGs (fold-change ≥ 1.2 ; adjusted p < 0.05) in splenic CD45.2^+^NGFR^+^ cells from mice at 4 weeks post-transplant. (F) Gene Set Enrichment Analysis (GSEA) plots of Hallmark MYC-target pathways. (G) Heatmap of DEGs from the MYC-target pathways. (H) MYC-GFP expression on CD45.2+NGFR+ cells from mice with percentage of MYC-GFP+ cells indicated (left), and average (± SD) GFP mean fluorescence intensity (MFI) (right). 2-way ANOVA: **, p < 0.01; ****, p < 0.0001; WT, n = 4; KO, n = 5 mice. (I) Flow cytometry analysis of CD34 versus MYC-GFP expression on CD45.2^+^NGFR^+^ cells with mean (± SD) frequencies of each gate in BM (top) and spleen (bottom). 2-way ANOVA: ****, p < 0.0001; WT, n = 4; KO, n = 5 mice.

To determine T-ALL burden in these animals, we enumerated leukemia cells in the bone marrow (BM), spleen, and peripheral blood by flow cytometry focusing on CD45.2^+^NGFR^+^ donor cells, which corresponded to ICN1-expressing cells. Unlike wildtype T-ALL mice that harbored large numbers of CD45.2^+^NGFR^+^ leukemic cells, KO T-ALL mice contained significantly fewer CD45.2^+^NGFR^+^ cells in BM, spleen and blood (**Figures 6D** and **S6E**). Furthermore, while these CD45.2^+^NGFR^+^ cells were phenotypically all CD4^+^CD8^+^ in WT T-ALL animals, the few KO CD45.2^+^NGFR^+^ cells had reduced frequencies of CD4^+^CD8^+^ cells and instead were also comprised of CD4^−^CD8^+^ and CD4^−^CD8^−^ fraction (**Figure S6F**), implying impaired ICN1-induced T-ALL lineage commitment. These results reveal an essential requirement for CHMP5 in ICN1-initiated T-ALL development *in vivo*.

To understand the molecular basis by which CHMP5 promoted T-ALL initiation by ICN1, we performed RNA-seq on sorted splenic CD45.2^+^NGFR^+^ (i.e., ICN1-transduced) cells from WT and KO T-ALL animals. Analysis of DEGs (fold-change ≥ 1.2; adjusted p < 0.05) between WT and KO-derived CD45.2^+^NGFR^+^ cells revealed 1157 upregulated genes and 706 downregulated genes. Similar to CHMP5-deficient human T-ALL cells above (**Figures 1A** and **1B**), unbiased pathway analysis of these DEGs also revealed marked downregulation of “MYC targets” in CD45.2^+^NGFR^+^ cells from KO T-ALL animals (**Figures 6E** and **6F**). Accordingly, CD45.2^+^NGFR^+^ cells from KO T-ALL animals expressed significantly less *Myc* and MYC target genes including *C1qbp, Ldha,* and *Phb2* (**Figure 6G** and **S6G)**.

To validate changes in MYC protein expression in our murine T-ALL model, we introduced a MYC-GFP fusion protein knock-in reporter allele^65^ into experimental donor mice, allowing us to quantify MYC protein expression by flow cytometry of GFP fluorescence in very few cells. Consistent with their *Myc* gene downregulation, *Chmp5*-deficient CD45.2^+^NGFR^+^ cells not only had fewer MYC(GFP)^+^ cells but also expressed significantly lower levels of MYC measured by GFP mean fluorescence intensity (MFI) protein (**Figure 6H**). These findings confirm that CHMP5 promoted the cellular machinery (namely, the BRD4-p300-MYC axis)^17–19^ that induces MYC gene and protein expression in this murine ICN1-driven T-ALL.

Murine and human T-ALL cells expressing CD34 and high levels of MYC have been shown to possess leukemia-initiating cell (LIC) potential^5,17,66,67^. We thus took advantage of the MYC-GFP reporter to assess the specific impact of CHMP5 deficiency on LICs in this ICN1-initiated murine T-ALL model. Remarkably, CD45.2^+^NGFR^+^ cells from KO T-ALL animals were not only depleted of MYC-GFP^hi^ CD34^+^ cells, but they also lacked any cells expressing CD34 in their BM and spleen (**Figures 6I**). Since MYC^hi^ and CD34^+^ T-ALL cells encompass a broad population of LICs in NOTCH1-driven T-ALL^5,17,66,67^, these data suggest that impaired T-ALL development in KO T-ALL animals likely resulted from abrogated LIC generation.

Collectively, our findings from this murine T-ALL model reveal a novel requirement for CHMP5 in T-ALL initiation and reinforced the CHMP5 dependence of oncogenic NOTCH1 in establishing and maintaining T-ALL transcriptional program exemplified by MYC. Notably, DEGs in ICN1-driven murine T-ALL cells and human NOTCH1-driven T-ALL cells lacking CHMP5 included multiple overlapping genes and pathways highlighted by *MYC* (and MYC target genes) (**Figures S6H** and **S6I**) which is induced by ICN1 by a BRD4-dependent mechanism^17–19^. Thus, regulation of the pro-leukemogenic transcriptional machinery in T-ALL cells by CHMP5 appears to be conserved in mice and human.

## Discussion

Using human and murine T-ALL models we have uncovered a critical role for the ESCRT protein CHMP5 in promoting the epigenetic and transcriptional program that enables T-ALL initiation and maintenance. The significance of CHMP5 function in T-ALL pathogenesis was supported by several lines of evidence: (i) oncogenic ICN1, which in wildtype BM progenitors initiated a lethal CD4^+^CD8^+^ T-ALL disease in mice, failed to cause T-ALL in BM progenitors where CHMP5 is selectively deleted in CD4-expressing thymocytes; (ii) CHMP5 deficiency in activated NOTCH1-driven human T-ALL impaired the T-ALL gene program exemplified by *MYC* and phenocopied the compromised metabolic fitness of MYC-deficient T-ALL cells; (iii) compared to normal T cells, CHMP5 is highly expressed across T-ALL subtypes and higher expression of CHMP5 correlated with worse T-ALL patient survival; (iv) CHMP5 deficiency synergistically improved chemotherapy efficacy, implicating CHMP5-driven mechanisms in chemoresistance in T-ALL cells. These effects of CHMP5 were at least in part driven by its ability to potentiate the p300-BRD4 interaction that promotes H3K27ac and Pol II pause release. Taken together, these findings highlight a previously unappreciated underlying mechanism in NOTCH1-driven T-ALL pathogenesis in which the ESCRT protein CHMP5 functions as critical positive regulator of the BRD4-p300 dependent transcription of T-ALL genes, including MYC that is essential for initiation and progression of these T-ALL disease subtype.

High MYC and CD34 expression define T-ALL cells with leukemia-initiating cell (LIC) properties in activated NOTCH1-driven T-ALL^5,17,66,67^. LIC activity is abolished in *Myc*-deficient T-ALL cells and MYC-GFP^lo/−^ murine T-ALL cells expressing the same MYC-GFP fusion reporter used in our study failed to cause disease in secondary recipients^17^. Thus, that CHMP5 deficiency resulted in the severe depletion of all CD34^+^ cells in primary murine T-ALL, including loss of the LIC-enriched CD34^+^MYC-GFP^hi^ fraction, indicated that failure of CHMP5-deficient progenitors to support ICN1-driven T-ALL was likely due to impaired generation of LICs. While we can’t exclude that deficiency in other genes besides *Myc* contributed to the loss of LIC-enriched cells in KO T-ALL mice, our data suggest a mechanism in which CHMP5 promoted T-ALL initiation at least in part through a BRD4-driven *Myc* transcriptional program. Intriguingly, CHMP5 is dispensable for normal CD4^+^CD8^+^ thymocyte generation^30^ (that lack MYC) but required for MYC^+^CD4^+^CD8^+^ T-ALL cells, implying that oncogenic ICN1 activity created a dependency on CHMP5 as part of the transcriptional rewiring required for T-ALL initiation.

Whether and how ESCRT proteins contribute to tumorigenesis has become of significant interest given their function in membrane remodeling is required for many key cellular processes. Accordingly, studies implicating ESCRT proteins in cancer have largely attributed their activity to membrane repair leading to inhibition of cell death^68–71^. We show here however that CHMP5 did not control T-ALL cell death but instead promoted leukemia by enabling transcription of T-ALL genes. Indeed, a fundamental insight from our study is the identification of a nuclear, chromatin-bound fraction of CHMP5 which, like other ESCRT proteins, was considered to be cytosolic^24,25,27^. In line with its nuclear localization, we found and validated an N-terminal bipartite NLS in CHMP5 that appears to be highly conserved across jawed vertebrates in which it first appeared, suggesting that CHMP5’s nuclear function was acquired later in evolution. How cytosolic versus nuclear CHMP5 trafficking is regulated is presently unclear but the presence of an NLS in CHMP5 suggest that it is actively imported into the nucleus.

To our knowledge our study represents the first report of a nuclear fraction of CHMP5 in mammalian cells. Previous reports from a yeast-two-hybrid screen predicted ESCRT protein interaction with nuclear proteins^72^ and some ESCRT proteins have been reported to promote nuclear envelope remodeling^73–75^. Only ESCRT protein CHMP1A has been previously demonstrated to interact with chromatin binding proteins in the nucleus^58^. Importantly however, CHMP5 and CHMP1A appear to play distinct roles in the nucleus and in T-ALL pathogenesis. Whereas mammalian CHMP1A, which also contains a bipartite NLS, functioned to recruit Polycomb-group proteins to silence genes^58^, CHMP5 promoted transcription by facilitating BRD4 recruitment to p300-catalyzed histone hyperacetylation at enhancers and super-enhancers. Moreover, we found that *CHMP5* expression levels, but not *CHMP1A*, were prognostic in T-ALL patients.

Since human T-ALL cells expressed significantly more (>5-fold) CHMP5 proteins than normal T-cells, our results would suggest that one mechanism by which oncogenes in T-ALL might hijack the transcriptional machinery and selectively control transcriptional output is by increasing CHMP5 expression. In turn, the quantity of CHMP5 dictated the amount of target T-ALL gene outputs. Indeed, our biochemical and genomic studies elucidated a mechanism in which nuclear CHMP5 amplified p300 interaction with BRD4 and consequently, improved both p300 and BRD4 binding selectively at enhancer and super-enhancer elements at key T-ALL genes (**Figure S4C**). Because p300 catalyzes H3K27ac of *cis* regulatory enhancers^49–51^ and BRD4 promotes p300 HAT activity^53^, this p300-BRD4 interaction conceivably operates as a positive feed-forward loop leading to further enriched BRD4 occupancy at enhancers and super-enhancers. At least in T-ALL cells, our data indicate that this process is critically dependent on CHMP5.

Surprisingly, while its pro-tumorigenic role has been demonstrated in other hematological cancers, notably acute myeloid leukemia (AML), knowledge of the specific role of p300 in T-ALL pathogenesis is limited. For example, translocations and fusions of p300, or its partner CBP, cause AML disease^76,77^ and small molecule inhibitors of p300/CBP suppress AML progression^78–80^. That CHMP5 deficiency -which mitigated T-ALL disease initiation and maintenance-impaired p300 recruitment to BRD4 is consistent with a pro-tumorigenic role for p300 in T-ALL pathogenesis. Therefore, it would be of interest to evaluate p300 inhibitors as therapeutic targets against T-ALL, especially those with underlying activating NOTCH1 mutations that display BRD4-dependency.

In conclusion, our study has uncovered a critical role for the ESCRT protein CHMP5 in promoting BRD4-p300-dependent super-enhancer formation that facilitates Pol II pause release and transcription of T-ALL initiation and maintenance genes. These findings provide mechanistic rationale for targeted CHMP5 depletion as a potential strategy against T-ALL. Loss of CHMP5 significantly depleted LICs *in vivo* suggesting that therapeutic targeting of CHMP5 can achieve more durable T-ALL suppression. Interestingly, high CHMP5 expression has also been reported in hepatocellular carcinoma^81^, and AML^82,83^ where its loss correlated with increased apoptosis^84,85^.Whether CHMP5 also regulates AML, in which the BRD4-dependent *MYC* super-enhancer is also a therapeutic vulnerability^52,86^, and other cancers that display a p300-BRD4 dependency remains to be determined. Therefore, future studies to elucidate CHMP5 function in these malignancies have the potential to significantly advance our understanding of transcriptional addiction mechanisms in cancer cells.

## Acknowledgments

We thank A. Tikhonova for critical reading of the manuscript and helpful suggestions, M. Carroll (University of Pennsylvania) for providing primary human T-ALL samples, A. Ferrando for CUTLL1 cells, X. Chen for MYC expression plasmids, and W. Pear for KOPT-K1 and DND-41 cell lines. We thank all current and former members of our laboratory for helpful discussions. This work was supported by grants from the US Department of Defense CA180768-W81XWH1910306 (S.A.); National Cancer Institute (NCI) K22 CA218467 (S.A.); National Institute of Allergy and Infectious Disease R01 AI1143992 (S.A.); American Cancer Society RSG-19-025-01-DDC (S.A.) and the Intramural Research Program of the National Institutes of Health, National Cancer Institute, Center for Cancer Research ZIA BC 012135 (S.A.). L.Z. is supported by grants from the National Heart, Lung, and Blood Institute R01 HL103827 and NCI R01 CA222064. K.U-W. was supported in part by a Cell and Molecular Biology Training grant NIGMS T32 GM008056 to the Case Western Reserve University School of Medicine. We acknowledge use of the high-performance computational capabilities of the Biowulf Linux cluster at the NIH (http://biowulf.nih.gov). All schematic cartoons were created with BioRender.com.

## Author contributions

Conceptualization, K.U-W., and S.A.; methodology, K.U-W., and S.A.; software, S.R., R.W., Q.T., Q.C., and D.M.; formal analysis, K.U-W., B.N.D., D.S.S., and S.A.; investigation, K.U-W., B.N.D., and S.A.; resources, L.Z., A.T., and D.S.S.; writing – original and final draft, K.U-W., and S.A.; writing – review & editing, all authors reviewed, edited, or commented on the manuscript; visualization, K.U-W., S.R., and S.A.; supervision, S.A.; funding acquisition, S.A.

## Methods

### RESOURCE AVAILABILITY

#### Lead contact

Further information and requests for resources and reagents should be directed to and will be fulfilled by the lead contact, Stanley Adoro.

#### Materials availability

Materials generated in this study are available upon request to the lead contact with a completed Materials Transfer Agreement.

#### Data and code availability

All sequencing data has been deposited to the Gene Expression Omnibus database (GEO) and are publicly available under the super-series GSE244200 as of the date of publication. RNA-seq on control (CT) and CHMP5 shRNA-depleted (KD) CUTLL1 human T-ALL cells can be downloaded under GEO accession number GSE244198. ChIP-seq of BRD4, Pol II, and H3K27ac in CT and KD CUTLL1 cells can be downloaded under GEO number GSE244197. Murine RNA-seq on wildtype and *Chmp5*-deficient ICN1-transduced CD45.2^+^NGFR^+^ splenocytes can be downloaded under GEO number GSE244199. This paper does not report original code.

### EXPERIMENTAL MODEL AND STUDY PARTICIPANT DETAILS

#### Mice

Six to eight-week-old male or female B6.SJL-Ptprca Pepcb/BoyJ (B6.SJL), NOD.Cg-Prkdc^scid^ Il2rg^tm1Wjl^/SzJ (NSG), B6;129-Myc^tm1Slek^/J (MYC-GFP)^65^, and B6.Cg-Tg(Cd4-cre)1Cwi/BfluJ (*Cd4*-Cre)^87^ mice were purchased from Jackson Laboratory. *Chmp5*^fl/fl^ mice (with loxP-flanked exons 3-7 of *Chmp5*) have been previously described^33^. All mice were maintained in specific-pathogen-free facilities at Case Western Reserve University (Cleveland, OH) or at the NCI campus in Frederick, Maryland under Institutional Animal Care and Use Committee approved protocols.

#### Cell culture

HEK293T (ATCC) and Plat-E cells (Cell Biolabs #RV-101) were cultured in DMEM supplemented with 10% FBS, 2 mM l-glutamine, 1 mM sodium pyruvate, 10 mM HEPES, pH 7.4, 1 × MEM nonessential amino acids, 50 IU/ml penicillin, and 50 μg/ml streptomycin. CUTLL1 (gift from Adolfo Ferrando, Columbia University) and primary T-ALL samples were cultured in complete RPMI (RPMI 1640 supplemented with 2 mM l-glutamine, 1 mM sodium pyruvate, 10 mM HEPES, pH 7.4, 1 × MEM nonessential amino acids, 50 IU/ml penicillin, 50 μg/ml streptomycin, 55 μM β-mercaptoethanol) with 20% FBS. Jurkat, Loucy, MOLT-3, MOLT-4, CEM, and SUP-T1 were purchased from ATCC; KOPT-K1 and DND-41 cells were provided by W. Pear (University of Pennsylvania) and cultured in complete RPMI with 10% FBS. HSB-2 was purchased from Sigma Aldrich and cultured in complete IMDM with 10% FBS. Primary cells were cultured in complete RPMI with 20% FBS. All cell cultures were maintained at 37°C in 5% CO_2_ incubators.

#### Primary human samples

Normal human T-cells were isolated from peripheral blood collected from the Hematopoietic Biorepository & Cellular Therapy Core at Case Western Reserve University. PBMCs were separated from the blood using a Ficoll density gradient (GE Healthcare #17144002). CD4 or CD8 cells were isolated using negative selection Miltenyi Biotec kits. Primary T-ALL patient samples were generously provided by Martin Carroll at the Stem Cell and Xenograft Core at the University of Pennsylvania. Patient information is provided in **Table 1**. Patient samples were collected in correspondence with approved protocols.

#### Xenografts

Primary patient samples were thawed in 30ml IMDM with 2.5% fetal bovine serum (FBS) and 10 ug/ml DNase, filtered and washed in PBS. At least 1 million live cells were retro-orbitally injected into NSG mice. Mice were bled by the tail vein to monitor leukemia progression by human CD45 expression. T-ALL cells were isolated from the spleen and bone marrow of mice by negative selection with mouse CD45 microbeads (Miltenyi Biotec #130-052-301).

#### Bone marrow transduction and transplantation

Bones were processed with ACK and a mortar and pestle. Cells were then washed with complete RPMI and filtered through a 40μm strainer. Bone marrow (BM) from CD45.2 mice was subjected to Lineage depletion (Miltenyi Biotec #130-110-470). CD45.2 BM cells were plated at 1×10^6^ cells/ml in a 10 cm plate in 10 ml of complete StemPro-34 SFM (Thermo Fisher #10639011) containing recombinant mouse cytokines (20 ng/ml IL-3, 50 ng/ml IL-6, 10 ng/ml Flt3L and 50 ng/ml SCF, PeproTech), 2 mM L-glutamine, 50 IU/ml penicillin, 50 μg/ml streptomycin and 55 μM β-mercaptoethanol. After 24 hours, cells were infected with retroviral supernatants on Retronectin coated plates in the presence of 8μg/ml polybrene and centrifugation at 1,000xg for 2 hours at 25°C. After 48 hours, BM from CD45.1+ mice for hemogenic support were subjected to CD3 depletion (Miltenyi Biotec #130-094-973) and cells were harvested and washed with PBS. 1 million CD45.2 and 250,000 CD45.1 cells were injected retro-orbitally into lethally irradiated (1000 rads) CD45.1 mice. Mice were monitored for engraftment and disease by flow cytometry of the blood and euthanized according to IACUC protocols when mice became too sick (weight loss, hunching, piloerection, lethargy).

### METHOD DETAILS

#### Murine cell isolation

Bone marrow, thymus and spleen were harvested and processed by mechanical dissociation to obtain single cells. Blood was collected either from the tail vein or medial saphenous vein into K2 EDTA coated collection tubes (BD Biosciences #365974). Splenocytes, blood, and bone marrow cells were treated with ACK lysis buffer (Gibco #A1049201) to lyse red blood cells. Single-cell suspensions were filtered through 40-μm strainers and resuspended in complete RPMI (RPMI 1640 supplemented with 2 mM l-glutamine, 1 mM sodium pyruvate, 10 mM HEPES, pH 7.4, 1 × MEM nonessential amino acids, 50 IU/ml penicillin, 50 μg/ml streptomycin, 55 μM β-mercaptoethanol, and 10% FBS).

#### In vitro drug treatments and viability assay

For viability assays T-ALL cells were cultured for 3 or 4 days with the indicated concentrations of drugs and analyzed with the Vybrant MTT Cell Proliferation Assay Kit (Thermo #V13154). Cytarabine (AraC; #S1648), and JQ1 (#S7110) were purchased from Selleck Chemicals, and Compound E from Enzo Life Sciences (#ALX-270-415-C250). Where applicable, cells were treated with 1 μM Dexamethasone (Selleck Chemicals #S1322) for 16-18 hours.

#### Flow cytometry analysis and cell sorting

Single-cell suspensions were washed with flow-cytometry buffer (PBS, 2% FBS and 1 mM EDTA) and stained with fluorochrome-conjugated antibodies for 30 min at 4°C in the dark. After washing, cells were resuspended in a flow-cytometry buffer and DAPI (4′,6-diamidino-2-phenylindole, 0.5 μg/ml; ThermoFisher #D1306) solution to exclude dead cells. MitoSOX (M36008), MitoTracker (M46753), ER Tracker (E34250) were acquired from ThermoFisher and used as indicated in the manufacturers protocol. Click-iT Plus OPP Alexa Fluor 647 Protein Synthesis Assay kit (Invitrogen #C10458) was used as per manufacturer’s instructions. Annexin-V was stained using Annexin-V APC (Biolegend #640941) and Annexin-V Binding buffer (Biolegend #422201) incubated with 7-AAD (Tonbo Bioscience #13-6993-T500) for 20 minutes at RT. Cell cycle analysis was performed using the Click-iT Plus EdU Alexa Fluor 647 Flow Cytometry Assay Kit (Invitrogen #C10634). Cells were incubated for 2 hours with 10uM EdU and stained with LIVE/DEAD™ Fixable Violet Dead Cell Stain Kit (Invitrogen #L34955) before fixation. After the Click-chemistry was performed, samples were incubated with 7-AAD (BD Biosciences #559925) for 10 minutes before acquisition. Flow cytometry sorting was performed on a BD FACS Aria or Aria-SORP and cell analysis on a BD LSR Fortessa and BD LSR-II. Flow analysis was done using FlowJo software. The antibodies used are listed in **Table 2**.

**Table 2.**
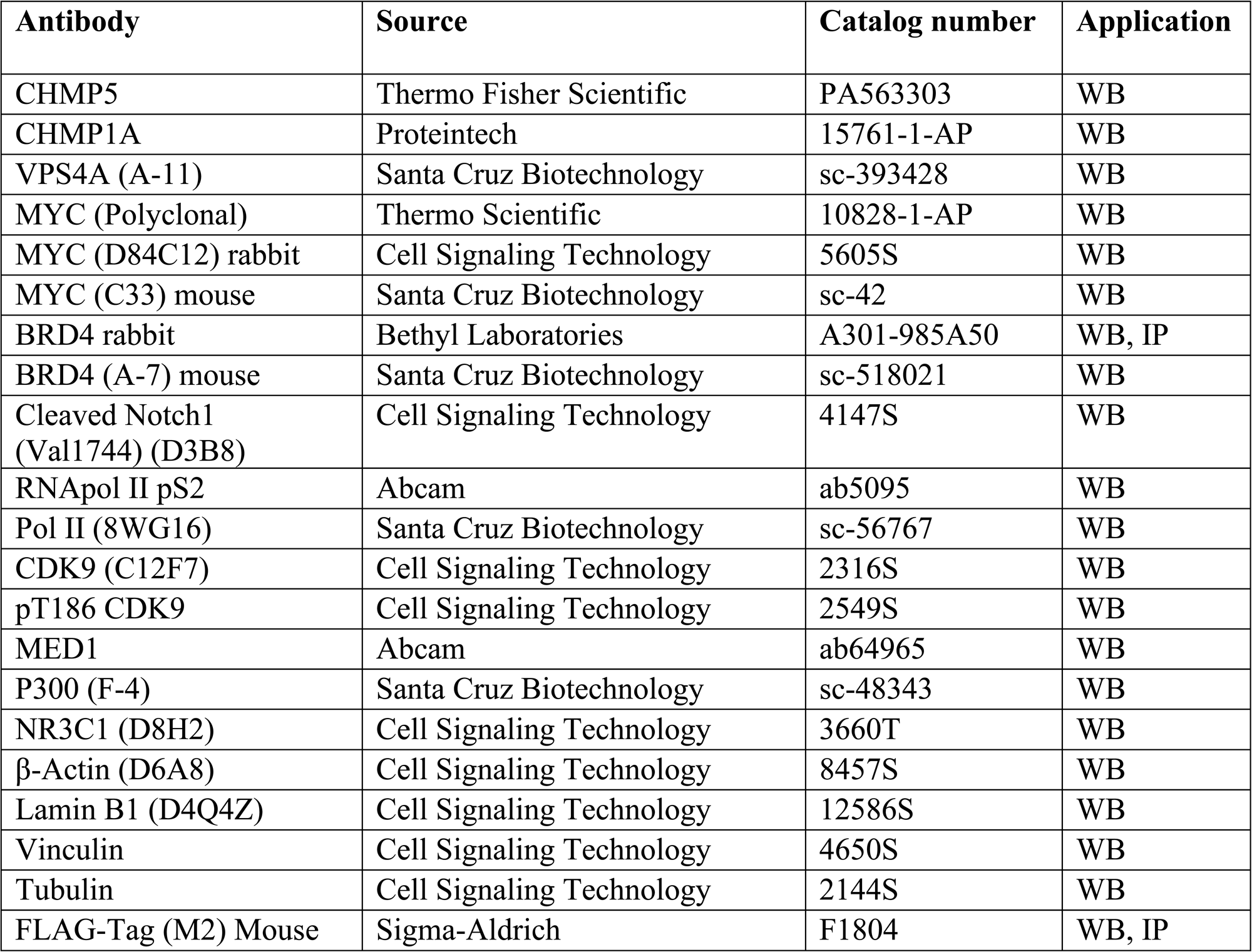
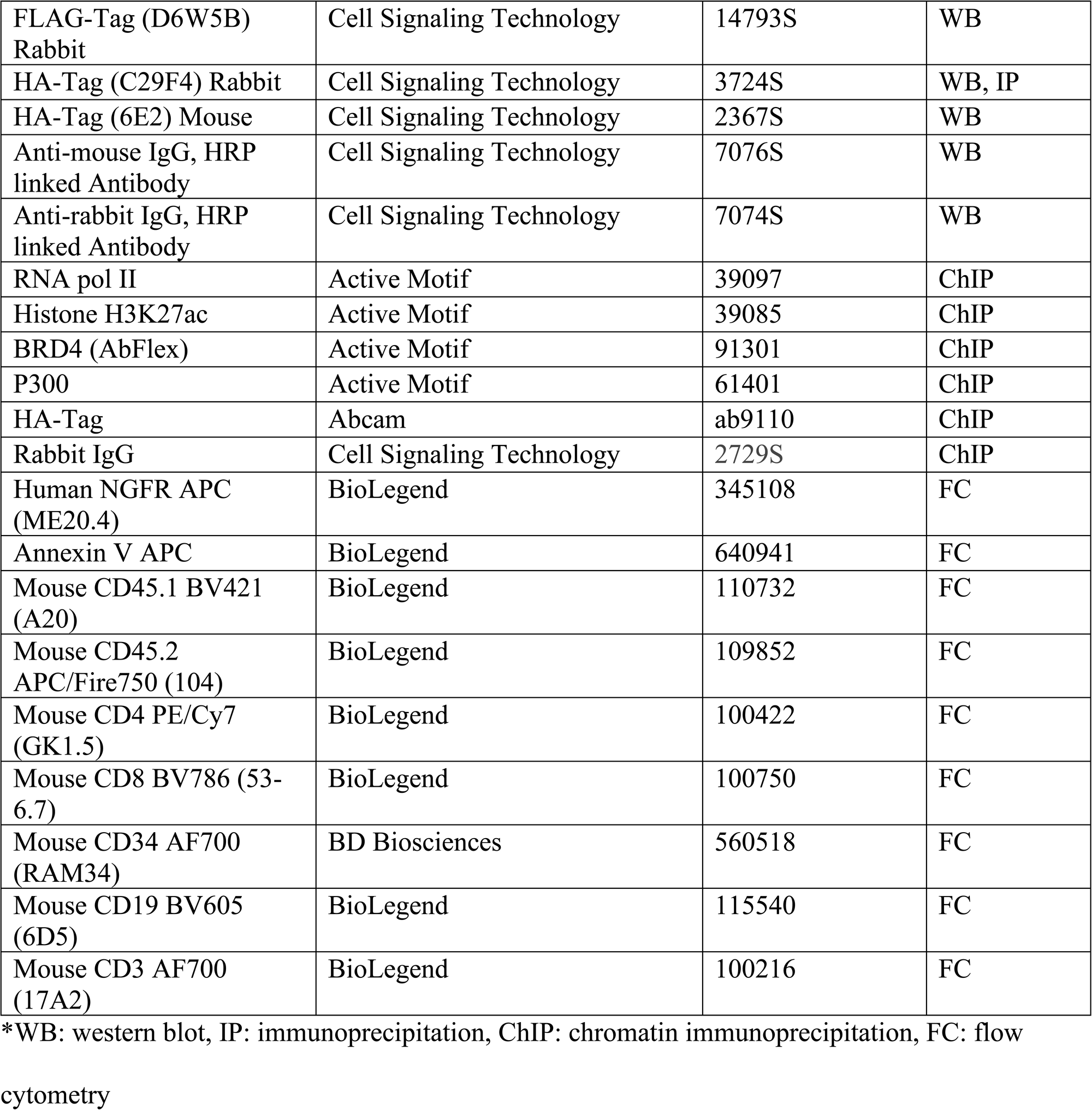
List of antibodies used for experiments.

#### Plasmids

C-terminal Flag-tagged CHMP5 (NP_084090.1) or N-terminal HA-tagged CHMP5 was amplified by PCR and cloned into pcDNA3.1 (Thermo #V79020) or pHAGE (Harvard) vectors. BRD4 overexpression plasmid for HEK293T transfection was purchased from Addgene (p6344 pcDNA4-TO-HA-Brd4FL #31351). shRNA constructs in a pLKO.1 vector targeting human CHMP5 were purchased from Sigma Aldrich along with 2 control non-targeting shRNAs. shRNAs targeting human MYC were a gift from Xi Chen (Baylor College of Medicine). shRNA sequences can be found in **Table 3**. ICN1-NGFR (pMIG) in which ICN1-transduced cells are marked by NGFR expression, was gifted to us by Lan Zhou (CWRU). The human CHMP5 delta NLS mutant was synthesized by Azenta Life Sciences.

**Table 3.**
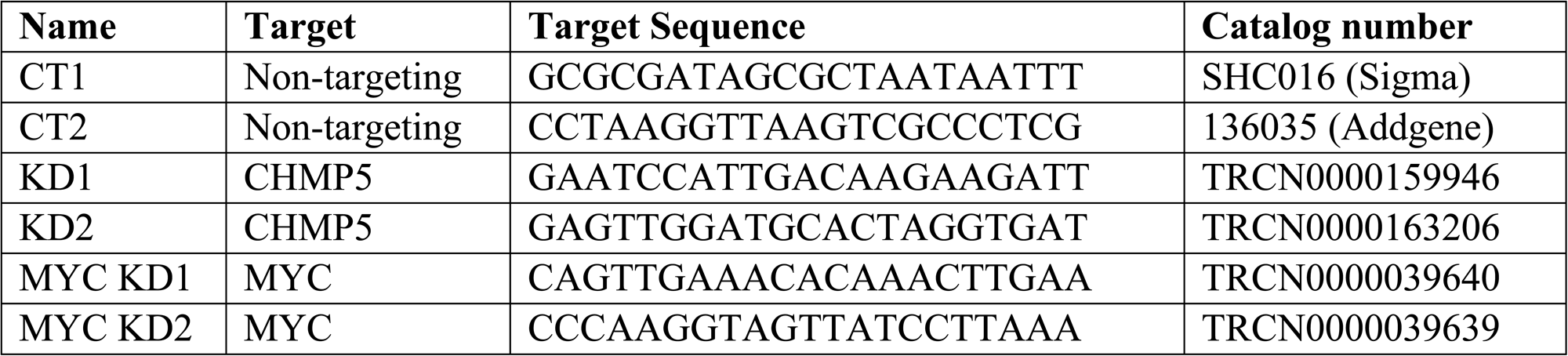
Target sequences for shRNAs.

#### Virus generation and cell transduction

Lentiviruses were generated by transfecting HEK293T cells with the plasmid of interest along with VSVg and delta 8.9 packaging plasmid vectors using Lipofectamine 3000 Transfection Reagent (Thermo #L3000075). After 48 hours, viral supernatants were harvested, filtered through a 0.45μm filter, concentrated with an Amicon Ultra-15 centrifugal filter (EMD Millipore #UFC903024) and stored at −80°C for future use.

Retroviruses were produced by transfecting Plat-E cells with the plasmid alone using Lipofectamine 3000. After 48 hours, viral supernatants were harvested and filtered, then concentrated using Retro-X concentrator overnight (Takara Bio #631455). Untreated tissue culture plates were coated with 25mg/ml Retronectin (Takara Bio #T100B) overnight at 4°C to be used for retroviral transduction. Plates were blocked the next day with 2% bovine serum albumin in PBS for 30 minutes at RT. Concentrated retrovirus was spun down on the RetroNectin coated plates for 2 hours before adding the cells. Cells were transduced with lentiviruses or retroviruses by adding 8μg/ml polybrene (Sigma Aldrich #TR-1003-G) and a previously determined amount of virus based on titers in a 6 well plate. The cells were then spun at 1000xg for 2 hours at room temperature and cultured for 48 hours.

For generation of shRNA knockdown cells, 48 hours after transduction with shRNA lentivirus, cells were selected with 1mg/ml puromycin (Invivogen #ant-pr-1) for 3-5 days. Deletion was validated by western blot and qPCR.

#### RNA isolation and quantitative real-time PCR

Total RNA was isolated from all cells using TRIzol reagent (Thermofisher, #15596026) and 100 to 500 ng of RNA was reverse transcribed using the high-capacity cDNA reverse transcription kit (Thermofisher, #4368814). Real time qPCR was done using Fast SYBR Green Master Mix (Applied Biosystems #43-856-12) and ran on QuantStudio six Flex real-time PCR system (Applied Biosystems # 4485691). Individual gene expression levels were calculated using the change in cycling threshold (ΔC_T_) method as 2^−ΔCT^, where ΔC_T_ is [C_T_ (gene of interest)-C_T_ (housekeeping gene)]. qPCR primer sequences are included in **Table 4**.

**Table 4.**
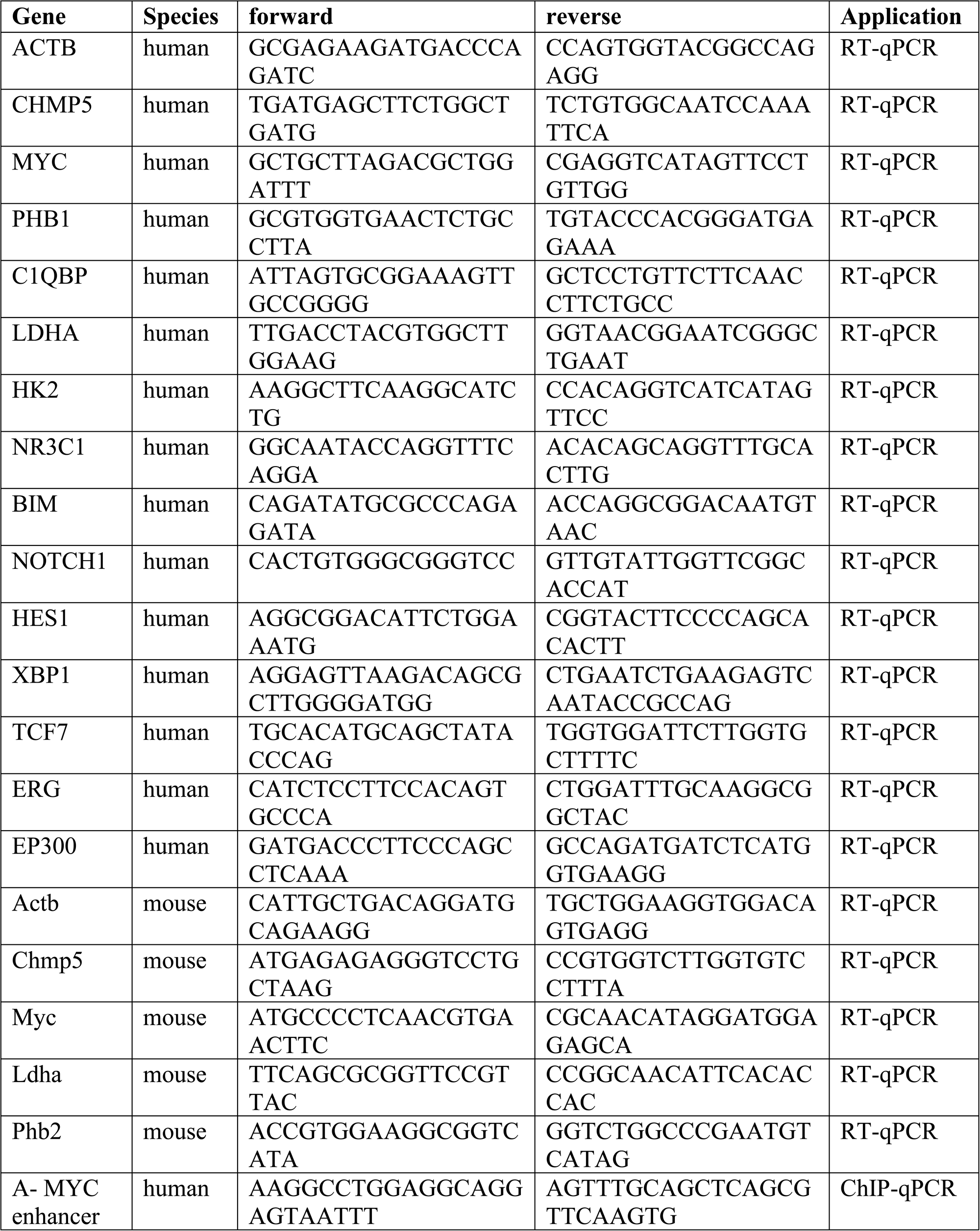
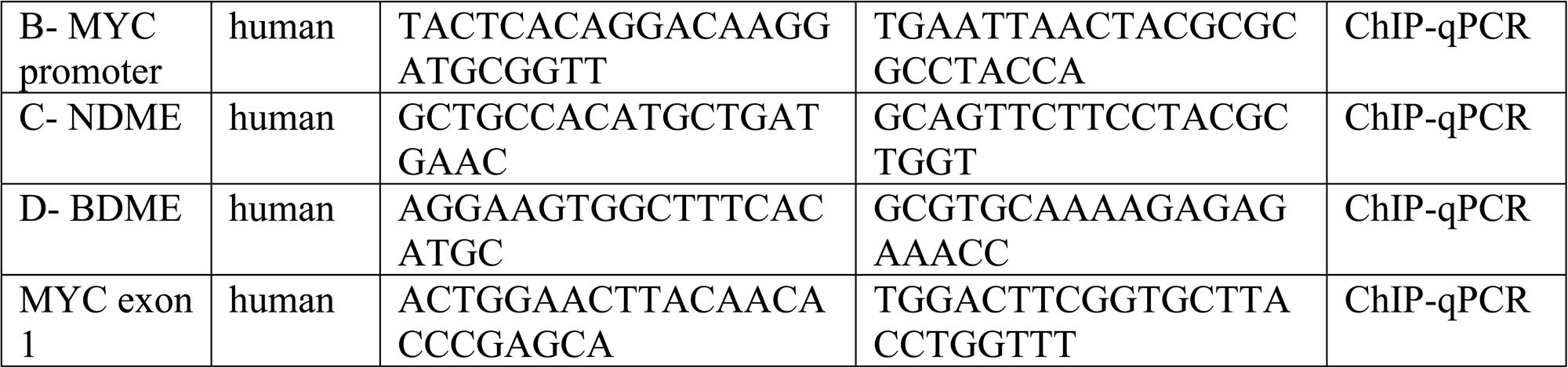
List of qPCR primers.

#### Immunoprecipitation and immunoblots

Cells are harvested and washed with ice-cold PBS by centrifuging at 5000rpm for 5 minutes at 4°C. Pellets were either stored at −80°C or lysed immediately; for western blots, cells were lysed with RIPA (25 mM Tris-HCl pH 7.6, 150 mM NaCl, 1% NP-40, 1% sodium deoxycholate, 0.1% SDS) buffer. For co-immunoprecipitations, cells were lysed with IP buffer (20 mM Tris-HCl, pH 8.0, 150 mM NaCl, 5 mM MgCl2, 0.5% NP-40). All lysis buffers were supplemented with 1X phosSTOP (Sigma Aldrich #4906837001) and 1X EDTA-free Protease Inhibitor Cocktail (Sigma Aldrich #4693159001). Lysates were collected after centrifuging cells at 13,000 rpm for 12 minutes at 4°C. Cell pellets used for fractionation were fractionated using the Nuclear Complex Co-IP kit from Active motif (#54001) according to their protocol.

Protein concentration was determined using Pierce™ BCA® Protein Assay Kits from Thermo fisher (#23225). 10-40 mg of protein was used for western blots, 200-500 mg of protein was used for immunoprecipitation. Immunoprecipitations were performed using magnetic Protein G Dynabeads (Thermo #10004D) with incubation of the primary antibody done overnight at 4°C with overnight rotation. NuPAGE LDS sample buffer (Thermo #NP0007) and Sample Reducing Agent (Thermo #NP0009) were added to all lysates and boiled at 70°C for 10 minutes. Samples were then run on NuPAGE 4-12% Bis-Tris Protein Gels (Thermo #NP0335BOX) and transferred to a PVDF membrane (EMD Millipore #IPVH00005). Blots were probed with primary antibodies overnight at 4°C in 5% blocking solution. After washing and 1 hour incubation with HRP-linked secondary antibodies, blots were developed using Pico- or Femto-chemiluminescent substrate (ThermoFisher) and visualized with autoradiography or digital imaging. Where indicated, band intensities were determined by densitometry using the ImageJ program (NIH).

Cell-free protein interaction assays were performed as previously described^11^. Recombinant CHMP5 was purchased from Abcam (ab134604), recombinant BRD4 was previously generated^11,88^, recombinant p300 was purchased from Active Motif (#81158). Recombinant proteins were diluted with Buffer D (20 mM HEPES, 100 mM KCl, 0.2 mM EDTA, and 20% vol/vol glycerol) to 0.1 mg/ml and 0.5 mg of each were incubated either individually or mixed in a 1:1 ratio in 500 ml of TBS (50 mM Tris-HCl, 150 mM NaCl) with protease inhibitors. Proteins were incubated overnight at 4°C with gentle rotation. Subsequently, 1ug of anti-FLAG (Sigma #F1804) or anti-BRD4 (Bethyl Laboratories #A301-985A50) was incubated with the proteins for 4 hours at 4°C with gentle rotation, and 40 ml of magnetic Protein G Dynabeads (Thermo #10004D) was added to each condition and incubated for 2 hours at 4°C with rotation. The beads were then washed 3 times with TBS+ 0.2% NP-40 and boiled at 70°C for 10 minutes in TBS with NuPAGE LDS sample buffer and reducing agent. The samples were run on NuPAGE 4-12% Bis-Tris Protein Gels. All antibodies used in these studies are listed in **Table 1**.

#### Protein stability assays

Cells were cultured in the presence of 20 μM MG132 (Sigma #M7449) for 6 hours unless indicated otherwise. For half-life experiments, cycloheximide (Sigma #01810) was used at 50 μg/ml. Cells were subsequently processed and subjected to immunoblotting as above. Protein expression was quantified by densitometry and half-life was calculated by normalizing protein levels during cycloheximide treatment to DMSO treated cells without cycloheximide.

#### Seahorse metabolic flux assays

Extracellular acidification rate (ECAR) was performed under glycolysis stress test and oxygen consumption rate (OCR) was performed under mitochondrial stress test. Briefly, 1 × 10^5^ CUTLL1 were plated per well on poly-D-lysine coated Seahorse XFe96 microplates (Agilent #101085-004) in XF RPMI medium (Agilent #103576-100) supplemented with 1mM Sodium Pyruvate, 2 mM L-glutamine, and 25 mM glucose for the Mito Stress Test (Agilent #103015-100), or 2 mM L-glutamine for the Glycolysis Stress Test (Agilent #103020-100). After an hour incubation at 37°C, OCR and ECAR were measured using a 96 well XF96 Extracellular Flux Analyzer (Agilent Technologies). Measurements were taken under basal conditions and following sequential addition of the drugs provided with the stress test kits. Fluoro-carbonyl cyanide phenylhydrazone (FCCP) was used at 1 mM final concentration. Basal ECAR was calculated from the samples in the Mito Stress Test medium containing glucose, and basal OCR was calculated from the samples in the Glycolysis Stress Test medium at the start of the assay.

#### Histology

Organs were fixed in 4% paraformaldehyde overnight and transferred to 70% ethanol before embedding. Embedding, sectioning and H&E staining was done at the Cleveland Digestive Diseases Research Core Center. Blood smears were made using Wright-Giemsa stain (Sigma #WG16). Images were taken using an Olympus IX73 microscope.

#### CUTLL1 RNA-seq sample processing, library preparation, and sequencing

RNA was isolated from CUTLL1 cells independently transduced with either shControl or shCHMP5 in triplicate using the Qiagen RNeasy kit. RNA was submitted BGI (Hong Kong, China) for sample QC, library preparation, and sequencing. RNA was quantified and integrity was checked using Agilent 2100 Bioanalyzer. Library preparation was done using the NuGEN Trio RNA-seq library preparation kit (Redwood City, CA). Libraries were sequenced with Illumina HiSeq X Ten PE150bp sequencing and 30 million reads per sample.

#### Murine RNA-seq sample processing, library preparation, and sequencing

CD45.2^+^NGFR(ICN1)^+^ cells were sorted from the spleens of chimera mice at median survival time-point (4 weeks post-transplantation). RNA was isolated from these cells using the TRIzol reagent and submitted to Azenta Life Sciences LLC (South Plainfield, NJ, USA) for library preparation and Ultra-low RNA sequencing. Total RNA samples were quantified using Qubit 2.0 Fluorometer (Life Technologies) and RNA integrity was checked using Agilent TapeStation 4200 (Agilent Technologies). The SMARTSeq HT Ultra Low Input Kit was used for full-length cDNA synthesis and amplification (Clontech), and Illumina Nextera XT library was used for sequencing library preparation. Briefly, cDNA was fragmented, and adaptor was added using Transposase, followed by limited-cycle PCR to enrich and add index to the cDNA fragments. Sequencing libraries were validated using the Agilent TapeStation and quantified by using Qubit Fluorometer as well as by quantitative PCR (KAPA Biosystems). The sequencing libraries were multiplexed and clustered on a flowcell. After clustering, the flowcell was loaded on the Illumina NovaSeq 6000 instrument according to manufacturer’s instructions. The samples were sequenced using a 2×150 Paired End (PE) configuration.

#### RNA-seq data processing and gene set enrichment analysis

Sequence reads for human samples control (CT) and knock-down (KD) were aligned to GRCh38 reference sequence annotated with Gencode release 39 with Star aligner (v2.7.9a)^89^ with --sjdbOverhang 100 and default parameters. Mouse wildtype (WT) and knockout (KO) samples were aligned to GRCm39 with Gencode release M27 with Star aligner (v2.7.9a) with --sjdbOverhang 150 and default parameters. STAR aligned reads gene counts were calculated with htseq (v0.11.4)^90^ with default parameters and differential expression analysis performed with DEseq2 (v1.38.3)^91^ software. Read counts were filtered to retain genes with at least 10 counts in the smallest group. Gene set enrichment analysis (GSEA) for the RNA-seq data was performed with GSEA 4.3.2^92^ and MsigDB Hallmark gene set collection^93^. For the comparison of CHMP5-regulated genes in human and mouse T-ALL, genes were extracted for mapping using BioMart R^94^ and Ensembl release 105^95^. Some DEGs are lost during the overlapping of human and mouse gene IDs.

#### Chromatin Immunoprecipitation (ChIP)

Fixation, isolation of the chromatin, immunoprecipitation and DNA isolation for ChIP analysis was done using the SimpleChIP® Enzymatic Chromatin IP Kit (CST #9003) and ran according to its protocol. Sonication was performed for 15 seconds on, 45 seconds off for 15 minutes. 10 ug of chromatin was isolated with 4 ug of indicated antibodies overnight. Antibodies used in these studies are listed in **Table 1**. qPCR was performed with 1ul of each per primer and compared to 2% input sample. Library preparation and sequencing of ChIP samples was performed by Azenta Life Sciences on the Illumina with 2 x 150bp, ∼350M PE reads.

#### ChIP-seq library preparation and sequencing

ChIP DNA samples were quantified using Qubit 2.0 Fluorometer (Life Technologies) and the DNA integrity was checked with 4200 TapeStation (Agilent Technologies). ChIP-Seq library preparation and sequencing reactions were conducted at Azenta US, Inc. (South Plainfield, NJ, USA). NEB NextUltra DNA Library Preparation kit was used following the manufacturer’s recommendations (Illumina). Briefly, the ChIP DNA was end repaired and adapters were ligated after adenylation of the 3’ends. Adapter-ligated DNA was size selected, followed by clean up, and limited cycle PCR enrichment. The ChIP library was validated using Agilent TapeStation and quantified using Qubit 2.0 Fluorometer as well as real time PCR (Applied Biosystems). The sequencing libraries were multiplexed and clustered on one lane of a flowcell. After clustering, the flowcell was loaded on the Illumina NovaSeq 6000 instrument according to manufacturer’s instructions (Illumina). Sequencing was performed using a 2×150 Paired End (PE) configuration. Image analysis and base calling were conducted by the Control Software (NCS). Raw sequence data (.bcl files) generated from the Illumina instrument was converted into fastq files and de-multiplexed using Illumina’s bcl2fastq 2.17 software. One mis-match was allowed for index sequence identification.

#### ChIP-seq data processing

Sequencing reads from BRD4, RNA Pol II, and H3K27 acetylation ChIP from control (CT) and knockdown (KD) and the input read samples were aligned to the reference genome UCSC hg38 with Bowtie2 (v2.4.5)^90^ with –local setting and unique reads were kept by filtering out unmapped, duplicated and multimapped reads with sambamba 0.8.2^96^ custom filters -F “[XS] == null and not unmapped and not duplicate”. MACS2 (v2.7.1)^97^ with default parameters was used for peak calling over the corresponding background. MACS2 peaks were annotated with ChIPseeker (v1.34.1) with TSS defined from −1kb to +1kb^98^. Heatmaps and metaplots displaying ChIP-seq occupancy were created with ngsplot^99^. ChIP tracks were generated using UCSC genome browser. The ROSE software^12,45^ was used to map typical and super enhancers with a maximum linking distance of 1 Mb for stitching.

#### Traveling Ratio (TR) calculation

Traveling Ratio (Pausing Index) calculated based on RNA Pol II ChIP-seq utilizing the PIC software (https://github.com/MiMiroot/PIC) as previously described in^43^ with default settings, – longest option to get the longest isoform, and gene body defined as +300bp to the end of annotated gene. Genes <1kb were removed from the analysis. Traveling Ratio (TR) was calculated for all genes, activated genes (promoters with H3K27ac peaks) as defined before, non-activated genes (no H3K27ac peak), BRD4 target (genes bound by BRD4 at promoters from ChIP-seq), and MYC target genes (HALLMARK MYC_TARGETS _V1 and V2). Traveling Ratio comparison between CT and KD samples was performed on log2 transformed ratios in R^100^ using Welch two sample t-Test.

#### Patient survival curves

RNA-seq and clinical data from TARGET-ALL-P2 cohort 1 was downloaded from the GDC data portal. Patients were divided into two groups according to the expression level of *CHMP5, VPS4A,* or *CHMP1A*; the top 20% were classified as highly-expressed group, and the bottom 20% were classified as lowly-expressed group. The entire group is comprised of the lowly-expressed group and highly-expressed group. Survival analysis is performed on these groups and survival curves are generated by using R-package ‘Survival’ and ‘Survminer’.

### QUANTIFICATION AND STATISTICAL ANALYSIS

Statistical details for experiments can be found in figure legends. Statistical analysis was performed using GraphPad Prism version 9.0. Except otherwise indicated, comparison between groups was determined with a two-tailed, unpaired Student’s t-test and differences were considered significant if p < 0.05 as follows: *, p = 0.01-0.05; **, p = 0.001-0.01; ***, p = 0.0001-0.001; ****, p < 0.0001. No statistical tests were used to predetermine sample size and no exclusion of data points was used.

---

**Figure S1.**
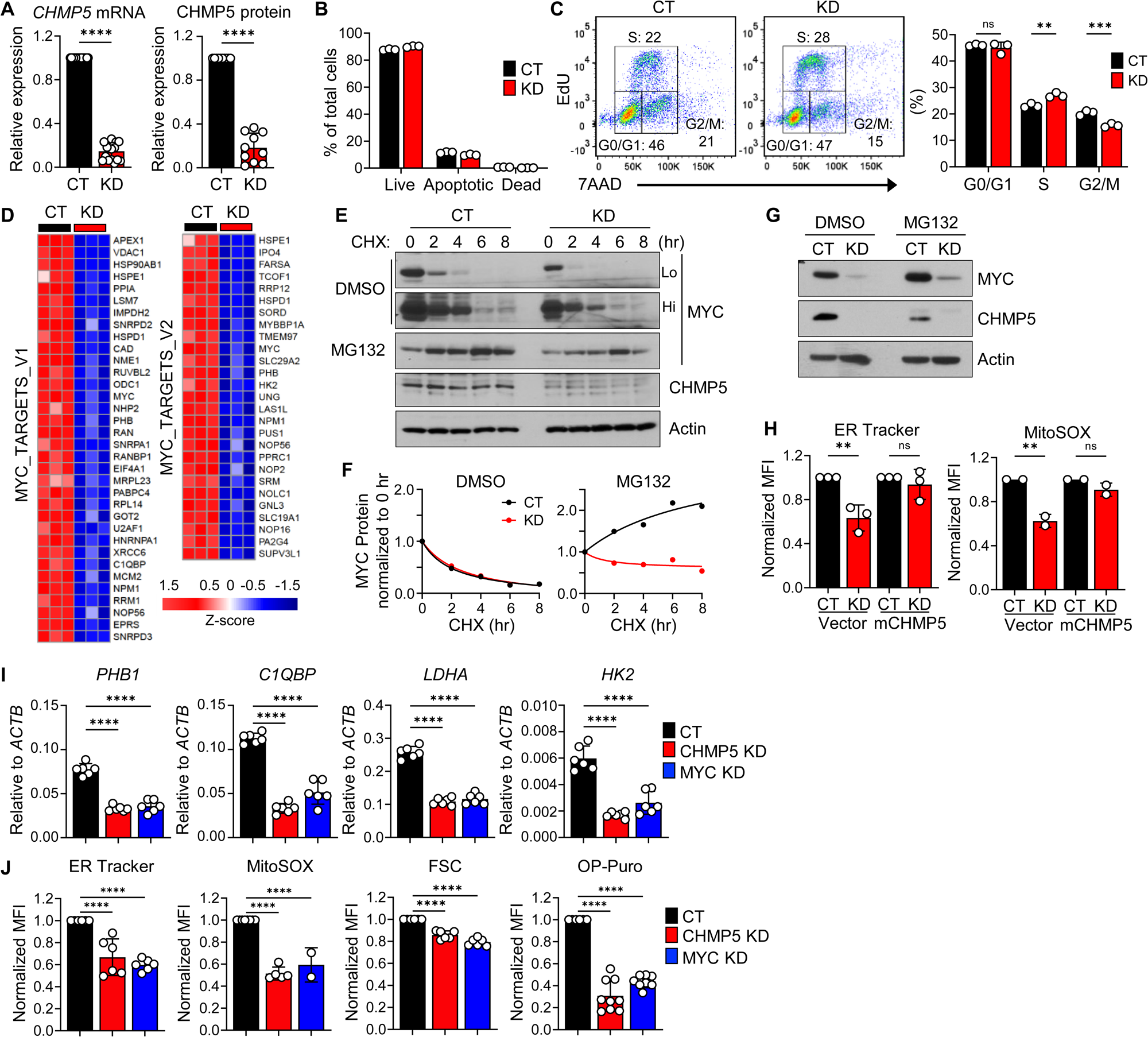
CHMP5 promotes the T-ALL gene program, related to Figure 1. (A) CHMP5 mRNA and protein expression relative to Actin and normalized to CT CUTLL1 cells. Data are presented as average (± SD) of replicates pooled from 5 independent experiments. Student’s t-test: ****, p < 0.0001. (B) Viability of CT and KD CUTLL1 cells determined by Annexin-V and 7-AAD staining. Data points are technical replicates and representative of 2 experiments. Difference between CT and KD are not significant. (p > 0.05). (C) EdU staining of CT and KD CUTLL1 cells with quantification of cells in different phases of cell cycle. Data are technical replicates and representative of 2 independent experiments. 2-way ANOVA: **, p < 0.01, ***, p < 0.001. (D) Heatmaps of DEGs from MYC-target gene pathways. (E) Western blot of CUTLL1 cells treated with cycloheximide (CHX) overtime with vehicle (DMSO) or 10 μM MG132. Lo, short exposure for MYC; Hi, long exposure for MYC. (F) Quantification of MYC expression in DMSO and MG132 treated samples from (**E**) overtime, normalized to the 0-time point of each group. Data is representative of 2 experiments. (G) Western blot of CT and KD CUTLL1 cells treated with vehicle (DMSO) or 10 μM MG132 for 6 hours. (H) Mean fluorescence intensity (MFI) for ER Tracker and MitoSOX dyes in CT and KD CUTLL1 transduced with murine CHMP5 (mCHMP5) or vector control. Data points are technical replicates. One-way ANOVA: **, p < 0.01; ns, p > 0.05 (I) mRNA expression of MYC-target metabolic genes. Data points are biological replicates from 3 independent experiments. One-way ANOVA: ****, p < 0.0001. (J) Flow analysis of CUTTL1 cells treated with shRNAs for CT, CHMP5 or MYC. MFIs are normalized to CT. Data points are biological replicates from 2-3 independent experiments. One-way ANOVA: ****, p < 0.0001.

**Figure S2.**
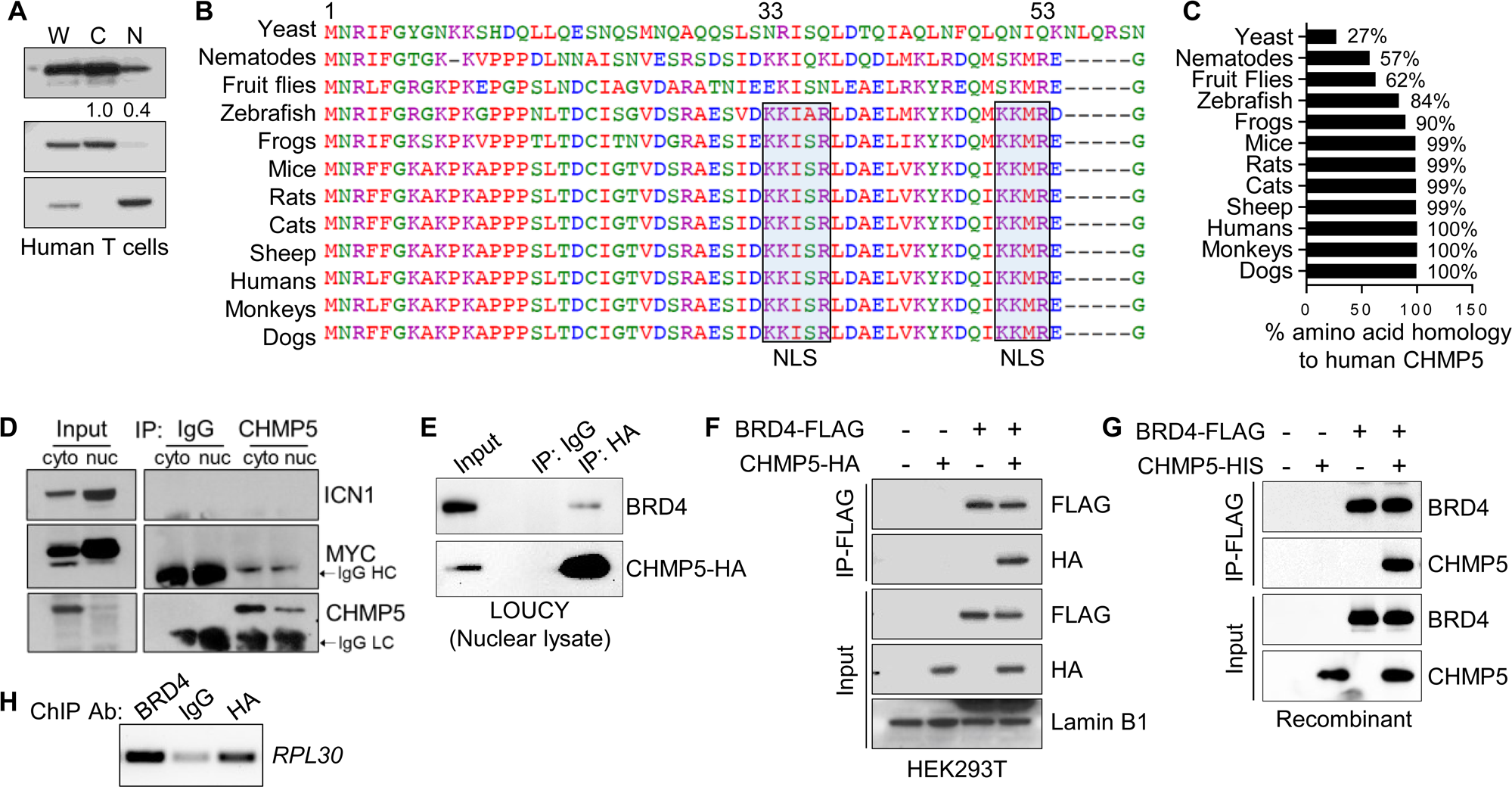
Identification of nuclear CHMP5-BRD4 interaction on chromatin, related to Figure 2. (A) Western blot of fractionated human T cells isolated from PBMCs. W, whole cell; C, cytoplasmic; and N, nuclear lysates. CHMP5 band intensity relative to the cytoplasmic band are indicated. (B) Clustal Omega alignment of the N-terminal amino acid sequences of CHMP5 from different species. Sequences corresponding to the bipartite nuclear localization sequence (NLS) are highlighted. (C) Amino acid sequence homology of CHMP5 from different species relative to human CHMP5. (D) Western blot of fractionated CUTLL1 cells transduced with CHMP5-HA and subjected to immunoprecipitation with isotype (IgG) or anti-HA antibodies and immunoblotted for MYC and ICN1. (E) Western blot of nuclear lysates in LOUCY cells subjected to immunoprecipitation with IgG or anti-BRD4 antibodies (F) Nuclear lysate of HEK293T cells co-transfected with BRD4-FLAG or CHMP5-HA and subsequently immunoprecipitated with anti-FLAG antibodies and immunoblotted with anti-HA or anti-FLAG antibodies. (G) Anti-FLAG immunoprecipitation of recombinant BRD4-FLAG and CHMP5-His recombinant protein. (H) PCR of *RPL30* on antibody (Ab) immunoprecipitated chromatin DNA from CUTLL1 cells transduced with CHMP5-HA run on a 1% agarose gel.

**Figure S3.**
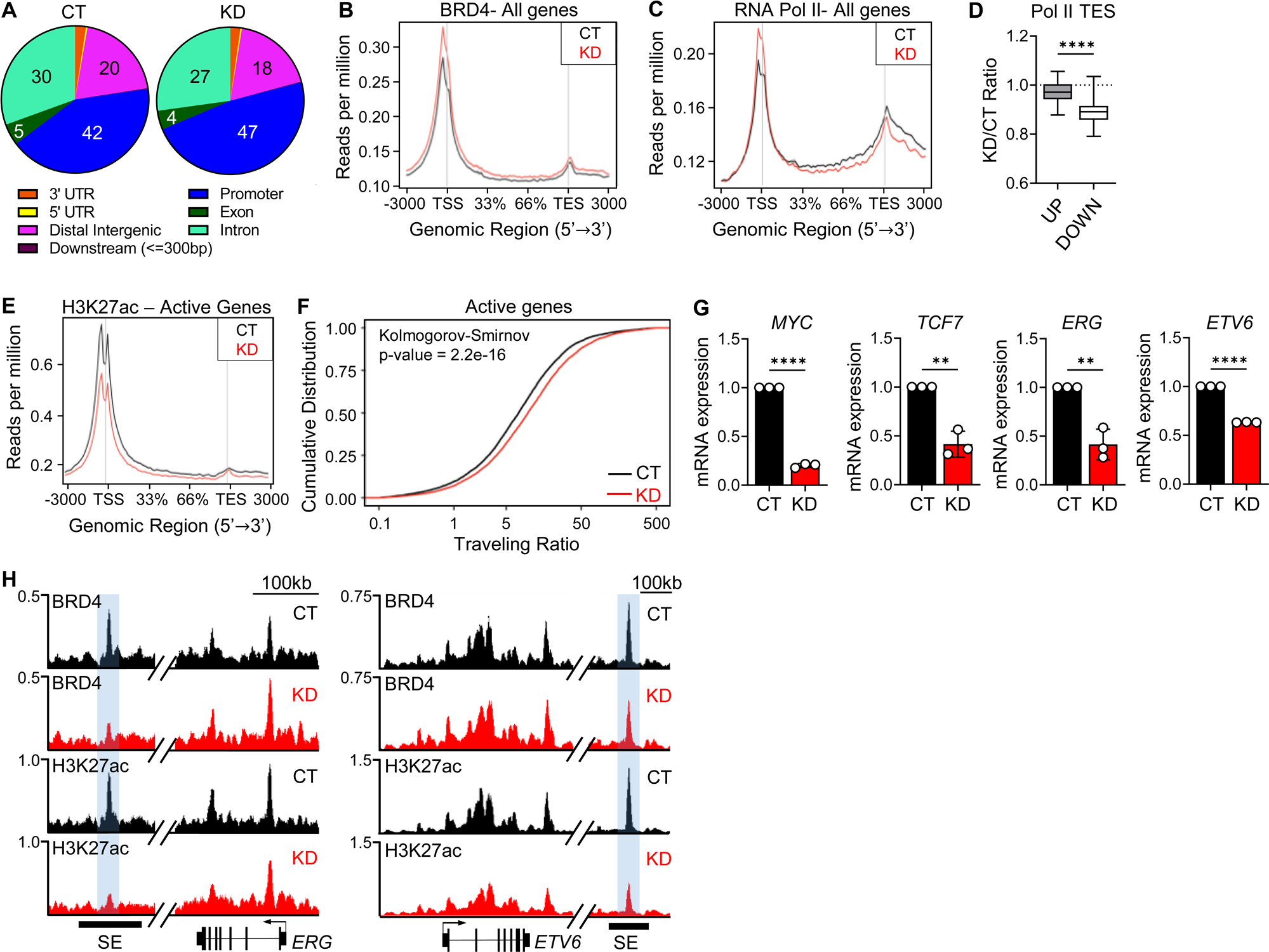
CHMP5 mediates BRD4-driven Pol II pause release and super enhancer formation, related to Figure 3. (A) Pie chart of genome-wide BRD4 occupancy in control (CT) and CHMP5-depleted (KD) CUTLL1 cells. (B) Metaplot of BRD4 binding at the TSS, gene body and TES across all genes in CT and KD CUTLL1 cells. (C) Metaplot of Pol II binding at the TSS, gene body and TES across all genes in CT and KD cells. (D) Box plot showing the ratio of KD to CT Pol II enrichment at the TES of UP and DOWN-regulated DEGs from RNA-seq on CT and KD CUTLL1 cells. Student’s t-test: ****, p < 0.0001. (E) Metaplot of H3K27ac density at the TSS, gene body and TES across active genes in CT and KD CUTLL1 cells. (F) Pol II traveling ratio at active genes (defined by H3K27ac signal at promoter). (G) Relative (normalized to control (CT) cells) mRNA expression of *MYC, TCF7, ERG,* and *ETV6* in CT and KD CUTLL1 cells. Data are average (± SD) of three biological replicates. Student’s t-test: **, p < 0.01; ****, p < 0.0001. (H) BRD4 and H3K27ac ChIP-seq tracks at the *ERG* and *ETV6* gene loci. SE, super-enhancer.

**Figure S4.**
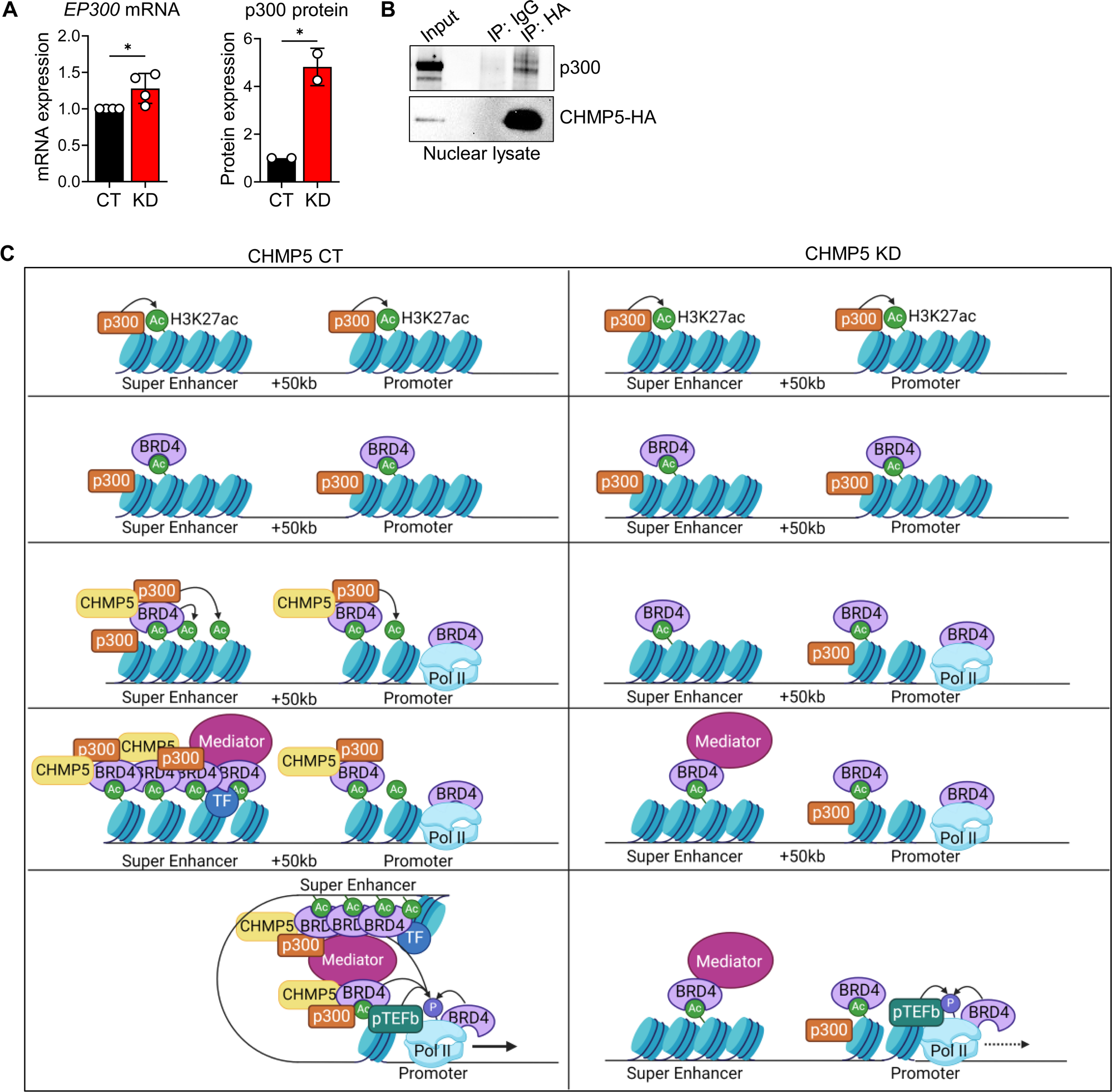
CHMP5 promotes the interaction between BRD4 and p300, related to Figure 4. (A) Relative mRNA (normalized to *ACTB*) and protein expression (normalized to Lamin B1) in CUTLL1 cells shown in **Figure 4A**. Data are average (± SD) of replicates from two independent experiments. Student’s t-test: *, p < 0.05. (B) Nuclear lysate from lentiviral CHMP5-HA-transduced CUTLL1 cells immunoprecipitated with isotype (IgG) or anti-HA antibody and immunoblotted for p300. (C) Mechanistic model of CHMP5-mediated regulation of epigenetic and transcriptional program in T-ALL cells. In wildtype T-ALL cells (left), CHMP5 potentiates the p300-BRD4 interaction that mediates H3K27 hyperacetylation of *cis* enhancers and super-enhancers. Subsequent assembly of core transcriptional factors at promoters and enhancers enables proximal-promoter and distal enhancer interaction that stimulate Pol II pause-release and transcriptional elongation of pro-leukemogenic genes. By contrast, CHMP5 deficiency (right) impairs the p300-BRD4 interaction, which reduces H3K27 acetylation, and disrupts super-enhancer formation and interaction with proximal-promoters leading to impaired transcription of T-ALL genes.

**Figure S5.**
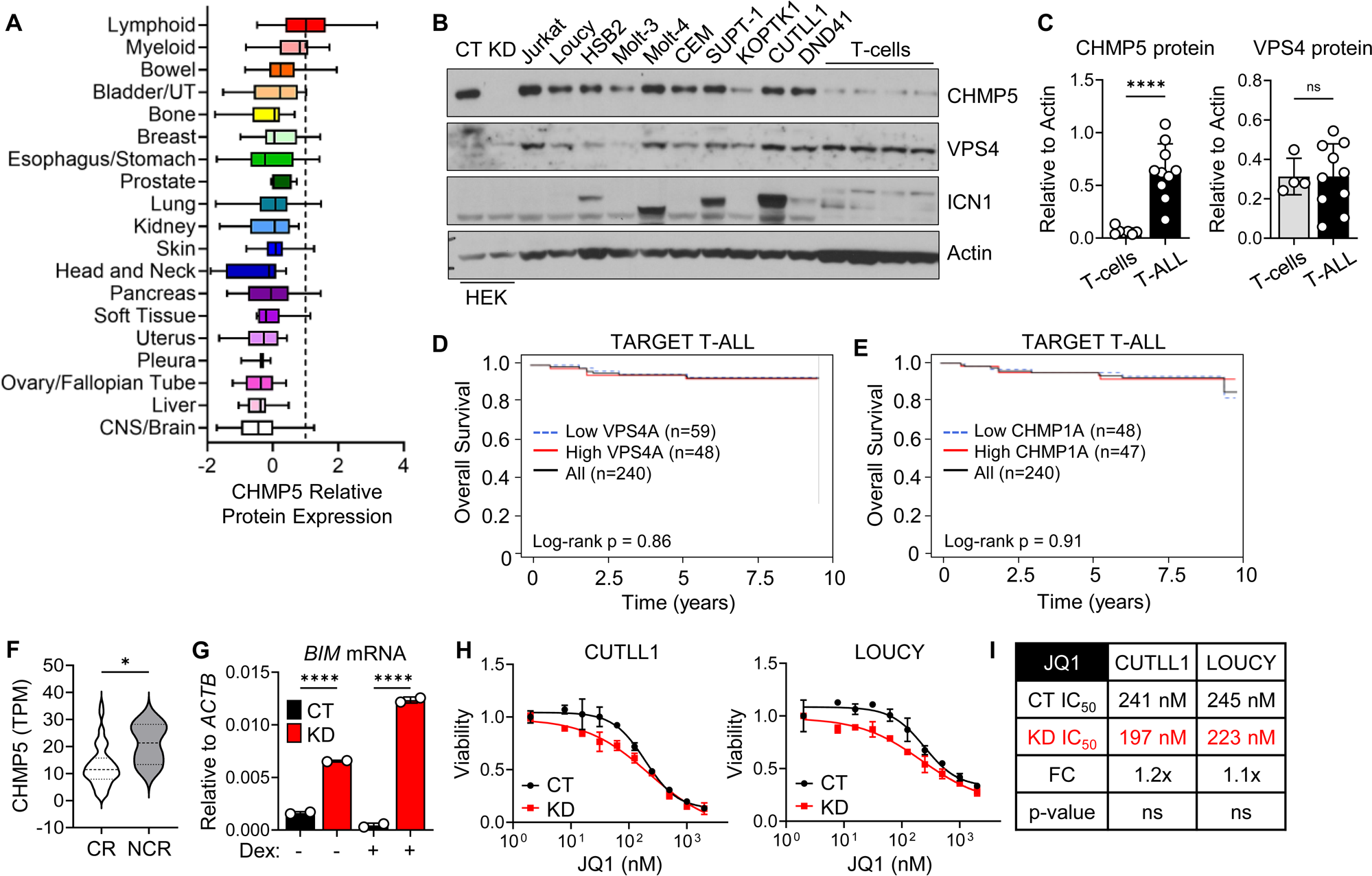
CHMP5 expression and impact on human T-ALL prognosis and chemoresistance, related to Figure 5. (A) Relative CHMP5 protein expression in cancer cell lines in order from highest to lowest average. Data from Cancer Cell Line Encyclopedia (CCLE). (B) Western blot of T-ALL cell lines and healthy donor human T-cells. Lysates from HEK293T (HEK) cells transduced with non-targeting shRNA (CT) or shCHMP5 (KD) lentivirus is used as control for anti-CHMP5 antibody specificity. (C) Quantification of CHMP5 and VPS4 protein relative to Actin from (B). Student’s t-test: ****, p < 0.0001. (D-E) Overall survival of pediatric T-ALL patients (TARGET T-ALL) expressing high (top 20%) and low (bottom 20%) levels of *VPS4A* (D) and *CHMP1A* (E). (F) mRNA expression of *CHMP5* in pediatric T-ALL patients that achieved complete remission (CR) (n = 65) or did not achieve complete remission (NCR) (n = 4). Student’s t-test: *, p < 0.05. (G) Expression of *BIM* in CUTLL1 cells from **Figure 6J**. Data are mean (± SD) of technical replicates. One-way ANOVA: ****, p < 0.0001 (H) Viability of CT and KD CUTLL1 and LOUCY treated with JQ1 for 3 days. Data are presented as mean (± SD) of 3 technical replicates, representative of 2 independent experiments. (I) IC_50_ for JQ1 in CUTLL1 and LOUCY cells. IC_50_ calculated by non-linear best-fit analysis. p-value calculated by 2-way ANOVA. FC, fold-change in CT versus KD IC_50_.

**Figure S6.**
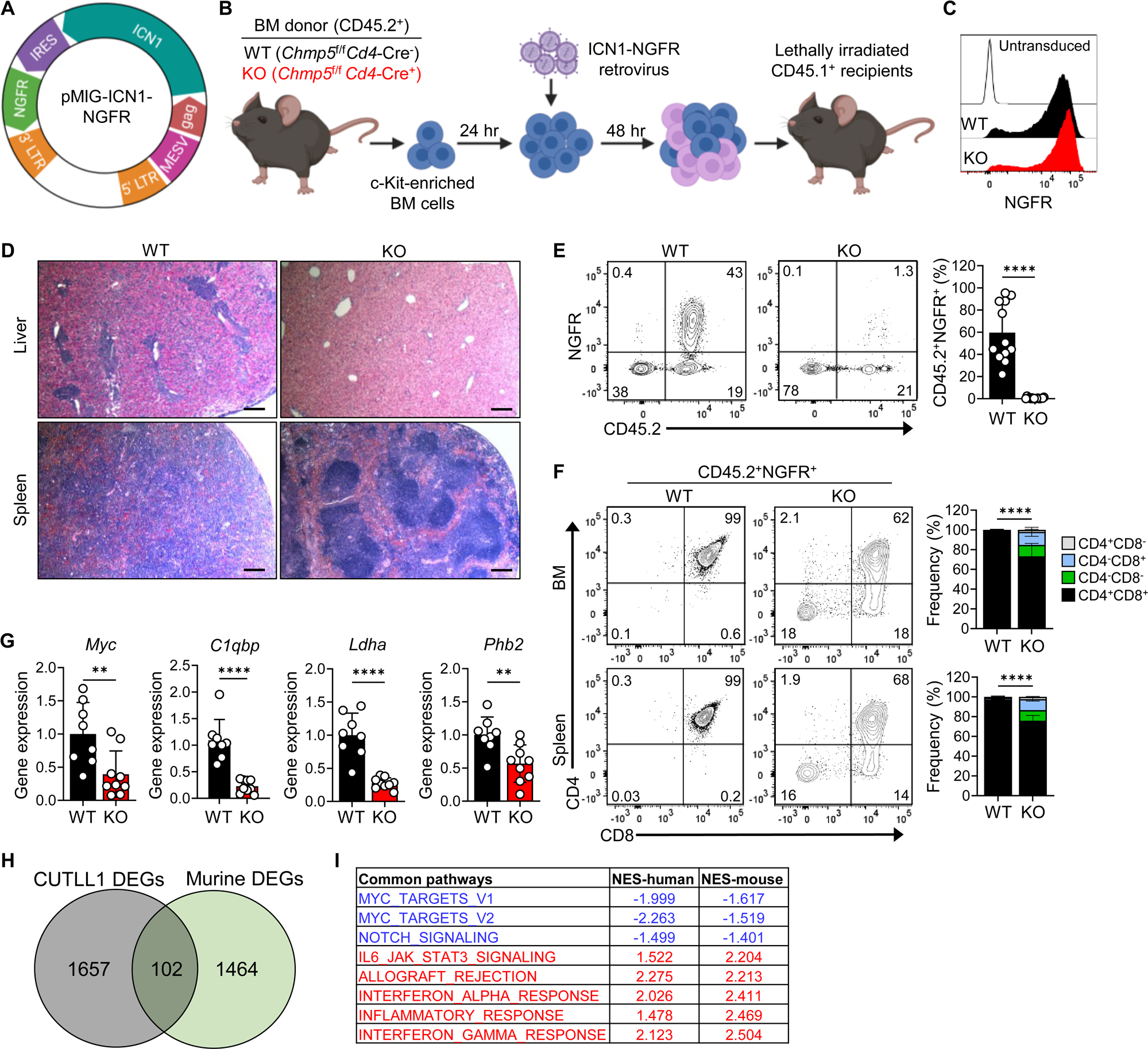
CHMP5 deficiency impairs T-ALL development and progression in vivo, related to Figure 6. (A) Plasmid map of bicistronic ICN1 and NGFR expression retroviral plasmid. IRES, internal ribosomal entry site. (B) Retrovirus-induced ICN1 leukemia mice experimental scheme. (C) NGFR expression 48 hours after transduction of WT and KO donor BM cells. (D) Hematoxylin and eosin staining of liver and spleen from leukemia mice. Scale bar = 200 µm (E) Representative flow cytometry analysis of blood from leukemia mice at 4 weeks post-transplant with average (± SD) frequency of CD45.2^+^NGFR^+^ cells shown in graph. Student’s t-test: ****, p < 0.0001; WT n=12, KO n=15 mice/group. (F) Flow cytometry plots of CD4 and CD8 expression on CD45.2^+^NGFR^+^ cells with average frequency (± SD) of gated subsets in WT n = 4 and KO n = 5 mice. 2-way ANOVA: ****, p < 0.0001; WT, n = 4; KO, n = 5 mice. (G) Average (± SD) mRNA expression of *Myc, C1qbp, Ldha,* and *Phb2* in splenic CD45.2^+^NGFR^+^ cells from WT (n = 8) and KO (n = 9) chimera mice. Expression is normalized to WT. Student’s t-test, **p < 0.01, ****p < 0.0001. (H) Venn-diagram of DEGs overlapping from CT and KD CUTLL1 (**Figure 1**), and WT and KO NGFR^+^ splenocytes. (I) Overlapping differentially expressed pathways from CT and KD CUTLL1, and WT and KO NGFR^+^ splenocytes.

## References

1 Weinstein, I. B. Addiction to Oncogenes--the Achilles Heal of Cancer. Science 297, 63–64 (2002). doi:10.1126/science.1073096

2 Bradner, J. E., Hnisz, D. & Young, R. A. Transcriptional Addiction in Cancer. Cell 168, 629–643 (2017). 10.1016/j.cell.2016.12.013

3 Delmore, J. E. et al. BET bromodomain inhibition as a therapeutic strategy to target c-Myc. Cell 146, 904–917 (2011). 10.1016/j.cell.2011.08.017

4 Zuber, J. et al. RNAi screen identifies Brd4 as a therapeutic target in acute myeloid leukaemia. Nature 478, 524–528 (2011). 10.1038/nature10334

5 Roderick, J. E. et al. c-Myc inhibition prevents leukemia initiation in mice and impairs the growth of relapsed and induction failure pediatric T-ALL cells. Blood 123, 1040–1050 (2014). 10.1182/blood-2013-08-522698

6 Wyce, A. et al. BET inhibition silences expression of MYCN and BCL2 and induces cytotoxicity in neuroblastoma tumor models. PLoS One 8, e72967 (2013). 10.1371/journal.pone.0072967

7 Patel, M. C. et al. BRD4 coordinates recruitment of pause release factor P-TEFb and the pausing complex NELF/DSIF to regulate transcription elongation of interferon-stimulated genes. Mol Cell Biol 33, 2497–2507 (2013). 10.1128/mcb.01180-12

8 Liu, W. et al. Brd4 and JMJD6-associated anti-pause enhancers in regulation of transcriptional pause release. Cell 155, 1581–1595 (2013). 10.1016/j.cell.2013.10.056

9 Anand, P. et al. BET bromodomains mediate transcriptional pause release in heart failure. Cell 154, 569–582 (2013). 10.1016/j.cell.2013.07.013

10 Devaiah, B. N. et al. BRD4 is a histone acetyltransferase that evicts nucleosomes from chromatin. Nature structural & molecular biology 23, 540–548 (2016). 10.1038/nsmb.3228

11 Devaiah, B. N. et al. BRD4 is an atypical kinase that phosphorylates Serine2 of the RNA Polymerase II carboxy-terminal domain. Proceedings of the National Academy of Sciences 109, 6927–6932 (2012). doi:10.1073/pnas.1120422109

12 Lovén, J. et al. Selective inhibition of tumor oncogenes by disruption of super-enhancers. Cell 153, 320–334 (2013). 10.1016/j.cell.2013.03.036

13 Creyghton, M. P. et al. Histone H3K27ac separates active from poised enhancers and predicts developmental state. Proc Natl Acad Sci U S A 107, 21931–21936 (2010). 10.1073/pnas.1016071107

14 You, M. J., Medeiros, L. J. & Hsi, E. D. T-Lymphoblastic Leukemia/Lymphoma. American Journal of Clinical Pathology 144, 411–422 (2015). 10.1309/ajcpmf03lvsblhpj

15 Weng, A. P. et al. Activating mutations of NOTCH1 in human T cell acute lymphoblastic leukemia. Science 306, 269–271 (2004). 10.1126/science.1102160

16 Herranz, D. et al. A NOTCH1-driven MYC enhancer promotes T cell development, transformation and acute lymphoblastic leukemia. Nat Med 20, 1130–1137 (2014). 10.1038/nm.3665

17 King, B. et al. The ubiquitin ligase FBXW7 modulates leukemia-initiating cell activity by regulating MYC stability. Cell 153, 1552–1566 (2013). 10.1016/j.cell.2013.05.041

18 Piya, S. et al. Targeting the NOTCH1-MYC-CD44 axis in leukemia-initiating cells in T-ALL. Leukemia 36, 1261–1273 (2022). 10.1038/s41375-022-01516-1

19 Sanchez-Martin, M. & Ferrando, A. The NOTCH1-MYC highway toward T-cell acute lymphoblastic leukemia. Blood 129, 1124–1133 (2017). 10.1182/blood-2016-09-692582

20 Knoechel, B. et al. An epigenetic mechanism of resistance to targeted therapy in T cell acute lymphoblastic leukemia. Nat Genet 46, 364–370 (2014). 10.1038/ng.2913

21 Yashiro-Ohtani, Y. et al. Long-range enhancer activity determines Myc sensitivity to Notch inhibitors in T cell leukemia. Proc Natl Acad Sci U S A 111, E4946–4953 (2014). 10.1073/pnas.1407079111

22 Kranz, A., Kinner, A. & Kölling, R. A family of small coiled-coil-forming proteins functioning at the late endosome in yeast. Mol Biol Cell 12, 711–723 (2001). 10.1091/mbc.12.3.711

23 Kohler, J. R. Mos10 (Vps60) is required for normal filament maturation in Saccharomyces cerevisiae. Mol Microbiol 49, 1267–1285 (2003). 10.1046/j.1365-2958.2003.03556.x

24 Babst, M., Katzmann, D. J., Estepa-Sabal, E. J., Meerloo, T. & Emr, S. D. Escrt-III: An endosome-associated heterooligomeric protein complex required for mvb sorting. Developmental Cell 3, 271–282 (2002). 10.1016/S1534-5807(02)00220-4

25 Azmi, I. F., Davies, B. A., Xiao, J., Babst, M., Xu, Z. & Katzmann, D. J. ESCRT-III family members stimulate Vps4 ATPase activity directly or via Vta1. Dev Cell 14, 50–61 (2008). 10.1016/j.devcel.2007.10.021

26 Liu, J., Kang, R. & Tang, D. ESCRT-III-mediated membrane repair in cell death and tumor resistance. Cancer Gene Ther 28, 1–4 (2021). 10.1038/s41417-020-0200-0

27 Vietri, M., Radulovic, M. & Stenmark, H. The many functions of ESCRTs. Nat Rev Mol Cell Biol 21, 25–42 (2020). 10.1038/s41580-019-0177-4

28 Alfred, V. & Vaccari, T. When membranes need an ESCRT: endosomal sorting and membrane remodelling in health and disease. Swiss Med Wkly 146, w14347 (2016). 10.4414/smw.2016.14347

29 Greenblatt, M. B. et al. CHMP5 controls bone turnover rates by dampening NF-κB activity in osteoclasts. J Exp Med 212, 1283–1301 (2015). 10.1084/jem.20150407

30 Adoro, S. et al. Post-translational control of T cell development by the ESCRT protein CHMP5. Nat Immunol 18, 780–790 (2017). 10.1038/ni.3764

31 Belver, L. & Ferrando, A. The genetics and mechanisms of T cell acute lymphoblastic leukaemia. Nat Rev Cancer 16, 494–507 (2016). 10.1038/nrc.2016.63

32 Palomero, T. et al. CUTLL1, a novel human T-cell lymphoma cell line with t(7;9) rearrangement, aberrant NOTCH1 activation and high sensitivity to gamma-secretase inhibitors. Leukemia 20, 1279–1287 (2006). 10.1038/sj.leu.2404258

33 Shim, J. H. et al. CHMP5 is essential for late endosome function and down-regulation of receptor signaling during mouse embryogenesis. The Journal of cell biology 172, 1045–1056 (2006). 10.1083/jcb.200509041

34 Gabay, M., Li, Y. & Felsher, D. W. MYC activation is a hallmark of cancer initiation and maintenance. Cold Spring Harb Perspect Med 4 (2014). 10.1101/cshperspect.a014241

35 Palomero, T. et al. NOTCH1 directly regulates c-MYC and activates a feed-forward-loop transcriptional network promoting leukemic cell growth. Proc Natl Acad Sci U S A 103, 18261–18266 (2006). 10.1073/pnas.0606108103

36 Lu, J. et al. Types of nuclear localization signals and mechanisms of protein import into the nucleus. Cell Communication and Signaling 19, 60 (2021). 10.1186/s12964-021-00741-y

37 Filippakopoulos, P. et al. Selective inhibition of BET bromodomains. Nature 468, 1067–1073 (2010). 10.1038/nature09504

38 Wang, H. et al. NOTCH1–RBPJ complexes drive target gene expression through dynamic interactions with superenhancers. Proceedings of the National Academy of Sciences 111, 705–710 (2014). doi:10.1073/pnas.1315023111

39 Yang, Z. et al. Recruitment of P-TEFb for stimulation of transcriptional elongation by the bromodomain protein Brd4. Mol Cell 19, 535–545 (2005). 10.1016/j.molcel.2005.06.029

40 Winter, G. E. et al. BET Bromodomain Proteins Function as Master Transcription Elongation Factors Independent of CDK9 Recruitment. Mol Cell 67, 5–18.e19 (2017). 10.1016/j.molcel.2017.06.004

41 Kanno, T. et al. BRD4 assists elongation of both coding and enhancer RNAs by interacting with acetylated histones. Nat Struct Mol Biol 21, 1047–1057 (2014). 10.1038/nsmb.2912

42 Uppal, S. et al. The Bromodomain Protein 4 Contributes to the Regulation of Alternative Splicing. Cell Rep 29, 2450–2460.e2455 (2019). 10.1016/j.celrep.2019.10.066

43 Chen, J. et al. VEGF amplifies transcription through ETS1 acetylation to enable angiogenesis. Nature Communications 8, 383 (2017). 10.1038/s41467-017-00405-x

44 Rahl, P. B. et al. c-Myc regulates transcriptional pause release. Cell 141, 432–445 (2010). 10.1016/j.cell.2010.03.030

45 Whyte, W. A. et al. Master transcription factors and mediator establish super-enhancers at key cell identity genes. Cell 153, 307–319 (2013). 10.1016/j.cell.2013.03.035

46 Hnisz, D. et al. Super-Enhancers in the Control of Cell Identity and Disease. Cell 155, 934–947 (2013). 10.1016/j.cell.2013.09.053

47 Chapuy, B. et al. Discovery and characterization of super-enhancer-associated dependencies in diffuse large B cell lymphoma. Cancer Cell 24, 777–790 (2013). 10.1016/j.ccr.2013.11.003

48 Bressin, A. et al. High-sensitive nascent transcript sequencing reveals BRD4-specific control of widespread enhancer and target gene transcription. Nat Commun 14, 4971 (2023). 10.1038/s41467-023-40633-y

49 Tie, F. et al. CBP-mediated acetylation of histone H3 lysine 27 antagonizes Drosophila Polycomb silencing. Development 136, 3131–3141 (2009). 10.1242/dev.037127

50 Weinert, B. T. et al. Time-Resolved Analysis Reveals Rapid Dynamics and Broad Scope of the CBP/p300 Acetylome. Cell 174, 231–244 e212 (2018). 10.1016/j.cell.2018.04.033

51 Narita, T. et al. Enhancers are activated by p300/CBP activity-dependent PIC assembly, RNAPII recruitment, and pause release. Mol Cell 81, 2166–2182 e2166 (2021). 10.1016/j.molcel.2021.03.008

52 Roe, J. S., Mercan, F., Rivera, K., Pappin, D. J. & Vakoc, C. R. BET Bromodomain Inhibition Suppresses the Function of Hematopoietic Transcription Factors in Acute Myeloid Leukemia. Mol Cell 58, 1028–1039 (2015). 10.1016/j.molcel.2015.04.011

53 Wu, T., Kamikawa, Y. F. & Donohoe, M. E. Brd4’s Bromodomains Mediate Histone H3 Acetylation and Chromatin Remodeling in Pluripotent Cells through P300 and Brg1. Cell Rep 25, 1756–1771 (2018). 10.1016/j.celrep.2018.10.003

54 Sanders, Y. Y. et al. Brd4-p300 inhibition downregulates Nox4 and accelerates lung fibrosis resolution in aged mice. JCI Insight 5 (2020). 10.1172/jci.insight.137127

55 Grosveld, F., van Staalduinen, J. & Stadhouders, R. Transcriptional Regulation by (Super)Enhancers: From Discovery to Mechanisms. Annu Rev Genomics Hum Genet 22, 127–146 (2021). 10.1146/annurev-genom-122220-093818

56 Van Vlierberghe, P. et al. ETV6 mutations in early immature human T cell leukemias. J Exp Med 208, 2571–2579 (2011). 10.1084/jem.20112239

57 Kourtis, N. et al. Oncogenic hijacking of the stress response machinery in T cell acute lymphoblastic leukemia. Nature Medicine 24, 1157–1166 (2018). 10.1038/s41591-018-0105-8

58 Stauffer, D. R., Howard, T. L., Nyun, T. & Hollenberg, S. M. CHMP1 is a novel nuclear matrix protein affecting chromatin structure and cell-cycle progression. Journal of Cell Science 114, 2383–2393 (2001). 10.1242/jcs.114.13.2383

59 Mullighan, C. G. The molecular genetic makeup of acute lymphoblastic leukemia. Hematology Am Soc Hematol Educ Program 2012, 389–396 (2012). 10.1182/asheducation-2012.1.389

60 Chen, B. et al. Identification of fusion genes and characterization of transcriptome features in T-cell acute lymphoblastic leukemia. Proceedings of the National Academy of Sciences 115, 373–378 (2018). doi:10.1073/pnas.1717125115

61 Real, P. J. et al. Gamma-secretase inhibitors reverse glucocorticoid resistance in T cell acute lymphoblastic leukemia. Nat Med 15, 50–58 (2009). 10.1038/nm.1900

62 Revollo, J. R., Oakley, R. H., Lu, N. Z., Kadmiel, M., Gandhavadi, M. & Cidlowski, J. A. HES1 Is a Master Regulator of Glucocorticoid Receptor–Dependent Gene Expression. Science Signaling 6, ra103–ra103 (2013). doi:10.1126/scisignal.2004389

63 Pui, J. C. et al. Notch1 expression in early lymphopoiesis influences B versus T lineage determination. Immunity 11, 299–308 (1999). 10.1016/s1074-7613(00)80105-3

64 Pear, W. S. et al. Exclusive development of T cell neoplasms in mice transplanted with bone marrow expressing activated Notch alleles. J Exp Med 183, 2283–2291 (1996).

65 Huang, C. Y., Bredemeyer, A. L., Walker, L. M., Bassing, C. H. & Sleckman, B. P. Dynamic regulation of c-Myc proto-oncogene expression during lymphocyte development revealed by a GFP-c-Myc knock-in mouse. Eur J Immunol 38, 342–349 (2008). 10.1002/eji.200737972

66 Gerby, B. et al. Expression of CD34 and CD7 on human T-cell acute lymphoblastic leukemia discriminates functionally heterogeneous cell populations. Leukemia 25, 1249–1258 (2011). 10.1038/leu.2011.93

67 Ma, W. et al. NOTCH1 signaling promotes human T-cell acute lymphoblastic leukemia initiating cell regeneration in supportive niches. PLoS One 7, e39725 (2012). 10.1371/journal.pone.0039725

68 Sadler, J. B. A. et al. A cancer-associated polymorphism in ESCRT-III disrupts the abscission checkpoint and promotes genome instability. Proc Natl Acad Sci U S A 115, E8900–e8908 (2018). 10.1073/pnas.1805504115

69 Dai, E., Meng, L., Kang, R., Wang, X. & Tang, D. ESCRT-III-dependent membrane repair blocks ferroptosis. Biochem Biophys Res Commun 522, 415–421 (2020). 10.1016/j.bbrc.2019.11.110

70 Neggers, J. E. et al. Synthetic Lethal Interaction between the ESCRT Paralog Enzymes VPS4A and VPS4B in Cancers Harboring Loss of Chromosome 18q or 16q. Cell Rep 33, 108493 (2020). 10.1016/j.celrep.2020.108493

71 Ritter, A. T. et al. ESCRT-mediated membrane repair protects tumor-derived cells against T cell attack. Science 376, 377–382 (2022). 10.1126/science.abl3855

72 Tsang, H. T. H., Connell, J. W., Brown, S. E., Thompson, A., Reid, E. & Sanderson, C. M. A systematic analysis of human CHMP protein interactions: Additional MIT domain-containing proteins bind to multiple components of the human ESCRT III complex. Genomics 88, 333–346 (2006). 10.1016/j.ygeno.2006.04.003

73 Olmos, Y., Perdrix-Rosell, A. & Carlton, J. G. Membrane Binding by CHMP7 Coordinates ESCRT-III-Dependent Nuclear Envelope Reformation. Curr Biol 26, 2635–2641 (2016). 10.1016/j.cub.2016.07.039

74 Willan, J. et al. ESCRT-III is necessary for the integrity of the nuclear envelope in micronuclei but is aberrant at ruptured micronuclear envelopes generating damage. Oncogenesis 8, 29 (2019). 10.1038/s41389-019-0136-0

75 Warecki, B., Ling, X., Bast, I. & Sullivan, W. ESCRT-III–mediated membrane fusion drives chromosome fragments through nuclear envelope channels. Journal of Cell Biology 219, e201905091 (2020). 10.1083/jcb.201905091

76 Iyer, N. G., Ozdag, H. & Caldas, C. p300/CBP and cancer. Oncogene 23, 4225–4231 (2004). 10.1038/sj.onc.1207118

77 Kitabayashi, I., Yokoyama, A., Shimizu, K. & Ohki, M. Interaction and functional cooperation of the leukemia-associated factors AML1 and p300 in myeloid cell differentiation. Embo j 17, 2994–3004 (1998). 10.1093/emboj/17.11.2994

78 Giotopoulos, G. et al. The epigenetic regulators CBP and p300 facilitate leukemogenesis and represent therapeutic targets in acute myeloid leukemia. Oncogene 35, 279–289 (2016). 10.1038/onc.2015.92

79 Gao, X. N. et al. A histone acetyltransferase p300 inhibitor C646 induces cell cycle arrest and apoptosis selectively in AML1-ETO-positive AML cells. PLoS One 8, e55481 (2013). 10.1371/journal.pone.0055481

80 Picaud, S. et al. Generation of a Selective Small Molecule Inhibitor of the CBP/p300 Bromodomain for Leukemia Therapy. Cancer Res 75, 5106–5119 (2015). 10.1158/0008-5472.Can-15-0236

81 Guo, Y., Shang, A., Wang, S. & Wang, M. Multidimensional Analysis of CHMP Family Members in Hepatocellular Carcinoma. Int J Gen Med 15, 2877–2894 (2022). 10.2147/ijgm.S350228

82 Wang, H. R. et al. PNAS-2: A Novel Gene Probably Participating in Leukemogenesis. Oncology 71, 423–429 (2006). 10.1159/000108576

83 Shahmoradgoli, M. et al. Antiapoptotic function of charged multivesicular body protein 5: A potentially relevant gene in acute myeloid leukemia. International Journal of Cancer 128, 2865–2871 (2011). 10.1002/ijc.25632

84 Wang, H. et al. The role of charged multivesicular body protein 5 in programmed cell death in leukemic cells. Acta Biochimica et Biophysica Sinica 45, 383–390 (2013). 10.1093/abbs/gmt028

85 Wang, H. R. et al. Anti-CHMP5 single chain variable fragment antibody retrovirus infection induces programmed cell death of AML leukemic cells in vitro. Acta Pharmacol Sin 33, 809–816 (2012). 10.1038/aps.2012.38

86 Shi, J. et al. Role of SWI/SNF in acute leukemia maintenance and enhancer-mediated Myc regulation. Genes Dev 27, 2648–2662 (2013). 10.1101/gad.232710.113

87 Lee, P. P. et al. A critical role for Dnmt1 and DNA methylation in T cell development, function, and survival. Immunity 15, 763–774 (2001). 10.1016/s1074-7613(01)00227-8

88 Maruyama, T. et al. A Mammalian bromodomain protein, brd4, interacts with replication factor C and inhibits progression to S phase. Mol Cell Biol 22, 6509–6520 (2002). 10.1128/mcb.22.18.6509-6520.2002

89 Dobin, A. et al. STAR: ultrafast universal RNA-seq aligner. Bioinformatics 29, 15–21 (2012). 10.1093/bioinformatics/bts635

90 Anders, S., Pyl, P. T. & Huber, W. HTSeq--a Python framework to work with high-throughput sequencing data. Bioinformatics 31, 166–169 (2015). 10.1093/bioinformatics/btu638

91 Love, M. I., Huber, W. & Anders, S. Moderated estimation of fold change and dispersion for RNA-seq data with DESeq2. Genome Biology 15, 550 (2014). 10.1186/s13059-014-0550-8

92 Subramanian, A. et al. Gene set enrichment analysis: a knowledge-based approach for interpreting genome-wide expression profiles. Proc Natl Acad Sci U S A 102, 15545–15550 (2005). 10.1073/pnas.0506580102

93 Liberzon, A., Subramanian, A., Pinchback, R., Thorvaldsdóttir, H., Tamayo, P. & Mesirov, J. P. Molecular signatures database (MSigDB) 3.0. Bioinformatics 27, 1739–1740 (2011). 10.1093/bioinformatics/btr260

94 Durinck, S., Spellman, P. T., Birney, E. & Huber, W. Mapping identifiers for the integration of genomic datasets with the R/Bioconductor package biomaRt. Nature Protocols 4, 1184–1191 (2009). 10.1038/nprot.2009.97

95 Cunningham, F. et al. Ensembl 2022. Nucleic Acids Research 50, D988–D995 (2021). 10.1093/nar/gkab1049

96 Tarasov, A., Vilella, A. J., Cuppen, E., Nijman, I. J. & Prins, P. Sambamba: fast processing of NGS alignment formats. Bioinformatics 31, 2032–2034 (2015). 10.1093/bioinformatics/btv098

97 Zhang, Y. et al. Model-based Analysis of ChIP-Seq (MACS). Genome Biology 9, R137 (2008). 10.1186/gb-2008-9-9-r137

98 Yu, G., Wang, L. G. & He, Q. Y. ChIPseeker: an R/Bioconductor package for ChIP peak annotation, comparison and visualization. Bioinformatics 31, 2382–2383 (2015). 10.1093/bioinformatics/btv145

99 Shen, L., Shao, N., Liu, X. & Nestler, E. ngs.plot: Quick mining and visualization of next-generation sequencing data by integrating genomic databases. BMC Genomics 15, 284 (2014). 10.1186/1471-2164-15-284

100 Team, R. C. (R Foundation for Statistical Computing, Vienna, 2022).

